# Molecular profiling of retinal pigment epithelial cell differentiation for therapeutic use

**DOI:** 10.1101/2021.01.31.429014

**Authors:** Sandra Petrus-Reurer, Alex R. Lederer, Laura Baqué-Vidal, Iyadh Douagi, Belinda Pannagel, Monica Aronsson, Hammurabi Bartuma, Magdalena Wagner, Helder André, Erik Sundström, Aparna Bhaduri, Arnold Kriegstein, Anders Kvanta, Gioele La Manno, Fredrik Lanner

## Abstract

Human embryonic stem cell-derived retinal pigment epithelial cells (hESC-RPE) are a promising cell source to treat age-related macular degeneration (AMD). Despite several ongoing clinical studies, detailed single cell mapping of the transient cellular and molecular dynamics from the pluripotent state to mature RPE has not been performed. Here we conduct single-cell transcriptomic analyses of 25,718 cells during differentiation as well as in embryonic and adult retina references, revealing differentiation progression through an unexpected initial cell diversification recapitulating early embryonic development before converging towards an RPE lineage. We also identified NCAM1 to track and capture an intermediate retinal progenitor with the potential to give rise to multiple neuroepithelial lineages. Finally, we profiled hESC-RPE cells after subretinal transplantation into the rabbit eye, uncovering robust *in vivo* maturation towards an adult state. Our detailed evaluation of hESC-RPE differentiation supports the development of safe and efficient pluripotent stem cell-based therapies for AMD.

## INTRODUCTION

The eye, by virtue of its accessibility and relatively isolated anatomical location, has emerged as a promising organ for gene and cell-based therapies to treat neurodegenerative diseases. A pathology that is particularly promising to tackle with these approaches is age-related macular degeneration (AMD), a major cause of severe vision loss affecting more than 180 million people globally (Gehrs et al., 2006). The dry form of the disease, for which no treatment is available, affects 80-90% of advanced patients and is characterized by well-demarcated areas of retinal pigment epithelium (RPE) loss and outer retinal degeneration (Ambati et al., 2003; Sunness, 1999). Human pluripotent stem cell (hPSC) derived RPE cells are thus of high interest for the development of cell replacement treatment options to halt disease progression, as currently being tested in several clinical trials (da Cruz et al., 2018; Kashani et al., 2018; Mandai et al., 2017; Schwartz et al., 2016; Song et al., 2015).

Notable efforts have been made towards developing strategies that ensure high purity RPE cell products using cell surface markers (Choudhary and Whiting, 2016; Plaza Reyes et al., 2020a). However, the focus on final product composition has often overshadowed the characterization of the intermediate stages appearing before a final steady state is reached. This gap has also been determined by the difficulty of deploying techniques that could systematically distinguish mixed phenotypes and off-target effects from cell heterogeneity and that would allow for a quantitative comparison to physiological references. In this perspective, the availability of single-cell RNA sequencing (scRNA-seq) represents a compelling opportunity.

scRNA-seq can systematically phenotype cell populations produced by differentiation protocols and its genome-wide readout is crucial to explore the unfolding of *in vitro* differentiation (Kulkarni et al., 2019; Kumar et al., 2017; Lederer and La Manno, 2020). For example, scRNA-seq can determine whether cells follow developmental or non-canonical paths to maturation (Cuomo et al., 2020; McCracken et al., 2020; Veres et al., 2019). Performing an unbiased analysis of the cell pool at intermediate stages might expose interesting relations between *in vitro* and *in vivo* processes and help to correctly identify potential risk sources for clinical translation (Begbie, 2013; Grove and Monuki, 2020; La Manno et al., 2016). Comprehensive single-cell atlases of embryonic and postnatal neurodevelopment are fundamental to assist in the evaluation of gene expression profiles measured *in vitro* (La Manno et al., 2016, 2020; Zeisel et al., 2018). Recent work has sought to decompose cellular heterogeneity of the embryonic and postnatal eye with scRNA-seq, but the similarity between the transient cell states arising in development and hPSC-derived intermediates en route to RPE lineage has not yet been evaluated (Collin et al., 2019; Cowan et al., 2020; Hu et al., 2019; Lo Giudice et al., 2019; Lukowski et al., 2019; Mao et al., 2019; Menon et al., 2019; Rheaume et al., 2018; Shekhar et al., 2016; Sridhar et al., 2020; Voigt et al., 2019). Importantly, both a reference-driven and an unbiased evaluation of the hPSC-RPE cell pool composition at different timepoints of the protocol are critical checkpoints for ensuring a safe and efficient RPE-based replacement therapy.

In this study, we performed scRNA-seq analyses during human embryonic stem cell (hESC) RPE differentiation using a directed and defined protocol established for clinical translation (Plaza Reyes et al., 2020a, 2020b). We demonstrate that the derived cells resemble embryonic retinal specification and eventually reach an RPE adult-like phenotype upon subretinal transplantation. These findings provide valuable insight into the developmental program of hESC-RPE differentiation and illustrate the required high quality of the derived cells to be used as future certified clinical product.

## RESULTS

### Human embryonic stem cells traverse gene expression space and sequentially mature into retinal pigment epithelium

To evaluate the process by which hESC-RPE are generated, we performed scRNA-seq throughout our established 60-day differentiation time course (Plaza Reyes et al., 2020a, 2020b). hESCs were differentiated on human recombinant laminin-521 (hrLN-521) using NutriStem hPSC XF medium to promote neuroepithelium induction. Activin A was provided at day 6 as a substitute for mesenchymal signaling to induce RPE fate (Cvekl and Wang, 2009; Fuhrmann et al., 2000; Fujimura, 2016). Cells were dissociated and replated at day 30 without Activin A, and maturation was completed by day 60 (**Figure 1A**). We profiled the culture at the undifferentiated starting state (hESC) and six consecutive time-points during the 60 days of differentiation (D7, D14, D30, D38, D45, D60). Morphological evaluation confirmed that changes in cell shape and size corresponded with the intended differentiation, as cells progressively assumed a more organized arrangement up until D60, where cultures appeared as a tight cobblestone monolayer of pigmented cells (**Figures 1B and S1A**).

**Figure 1.**
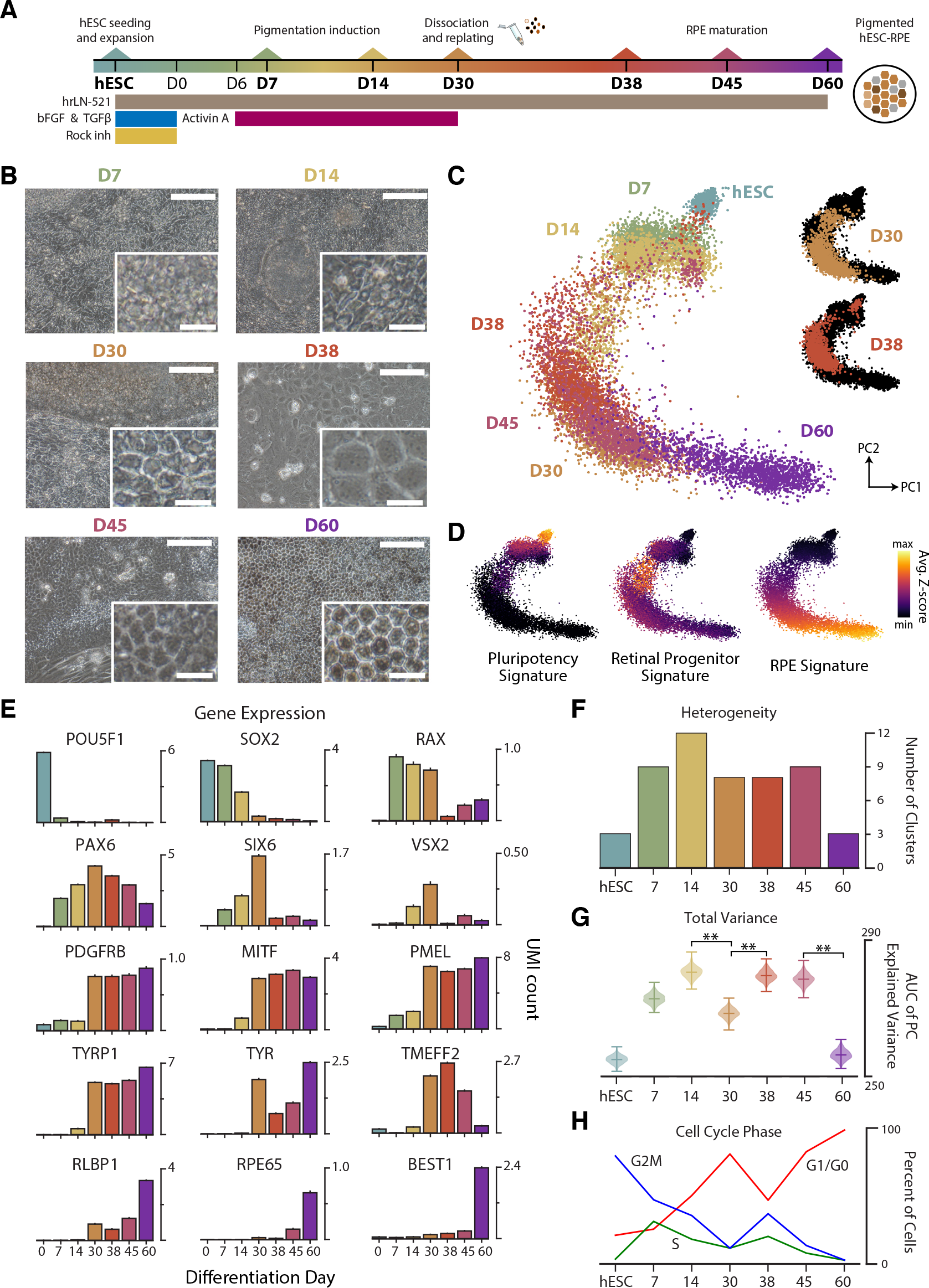
Global scRNA-seq characterization of the hESC-RPE differentiation time course. (A) Schematic of the hESC-RPE differentiation protocol. scR-NA-seq was performed at the seven bolded time points. (B) Brightfield images of hESC-RPE cultures. Scale bars: 100μm; inset 20μm. (C) Principal component representation of 11,791 single cells obtained using 42 well-characterized marker genes (6 pluripotency, 20 retinal progenitor, and 16 RPE genes). (D) Signature scores constructed from 42 marker genes (see Methods). (E) Bar charts indicating the average gene expression levels of pluripotent, retinal progenitor, and RPE cell markers at different time points. (F) Bar plot of the number of clusters identified. (G) Cumulative explained variance curve using area under the principal components for each timepoint, applied to estimate how much the variance accumulates over sets of correlated genes (i.e. biological-driven variability), as opposed to uniformly across genes (i.e. white noise). **p<0.01. (H) Line-plots reporting the percentage of cells assigned to the different cell cycle phases as deterimned by scRNA-seq data scoring. Cycling: S and G2/M; Growth: G1/G0. Bars represent mean +/-SEM from all cells at each time point. See also Figure S1.

We next applied prior biological knowledge to perform a global assessment of 11,791 single-cell transcriptomes and to verify overall population progression through the fundamental stages of RPE development. Using 42 marker genes for pluripotent, retinal progenitor, and RPE identities, we studied how cells traversed a reduced gene expression space. The visualization of these expression profiles using the principal components showed that, with time, cells progressively moved away from the pluripotent state towards a mature RPE identity (**Figures 1C, S1B, and S1C**). Gene signature scores detected a loss of the pluripotency signature (38-fold decrease in signature score from hESC to D30), an increased progenitor status at intermediate days (6.5-fold increase from hESC to D30), and a rise of mature RPE upon protocol conclusion (1.7-fold increase from D38 to D60) (**Figures 1D, S1D, and S1E**). Temporal assessment of gene expression confirmed a coherent sequence of expression waves, with pluripotency genes (*POU5F1, SOX2*) leading and being downregulated in favor of progenitor genes (*RAX, PAX6, SIX6, VSX2*), eventually trailed by early (*PDGFRB, MITF, PMEL, TYRP1, TYR, TMEFF2*) and late (*RLBP1, RPE65, BEST1*) maturation genes (Brandl et al., 2014; Schmitt et al., 2009; Sparrow et al., 2010) (**Figure 1E**). These analyses indicated that the differentiation protocol drives the cell pool towards RPE maturation through a path broadly consistent with the developmental process intended to be captured *in vitro*.

### Heterogeneity analysis reveals changes in cell diversity during RPE differentiation

Interestingly, we observed deviations from a uniform progression towards RPE. D30 cells appeared more differentiated towards an RPE fate than cells at D38, likely a response to dissociating and replating the cells at day 30 (**Figures 1C and S1C**). A subset of intermediate cells did not exhibit a strong signature for any of the three identities considered. This suggested a more complex and nonlinear differentiation process than anticipated as well as the presence of additional cell types not captured by our global analyses (**Figures 1D**).

Intrigued, we sought to harness the full phenotyping potential of scRNA-seq and achieve an in-depth description of heterogeneity at all stages. We applied balanced cell clustering and defined 52 clusters that collectively explained 84% of the variance (see Methods; **Figures 1F and S1F**). Initial (hESCs) and final (D60) samples had fewer clusters than intermediate time points as well as mutually exclusive and uniform expression of pluripotency and RPE genes, alluding to a diversity expansion during neuroepithelium induction (**Figures S1G and S1H**).

We next calculated how much variance accumulated in correlated gene modules as opposed to uncorrelated genes and interpreted this quantity as a measure of biological heterogeneity (see Methods; **Figure S1I**). Studying these values along with the number of clusters revealed that hESCs and D60 cells harbored a significantly lower heterogeneity compared to intermediate time points (**Figure 1G**). While a decrease of heterogeneity was detected from D14 to D30, suggesting an initial convergence towards RPE fate, we observed an increase from D30 to D38, hinting at an effect of tissue dissociation, replating, or Activin A removal on cell composition. This was consistent with proliferation trends: a decrease in cycling cells (S and G2/M phases) from hESC to D30, followed by an increased fraction from D30 to D38 and a second decrease from D38 to D60 (**Figure 1H**).

### Early hESC-RPE differentiation recapitulates the cellular diversity of the anterior neural tube and optic vesicle

To determine the identity of the 24 cell clusters found during pigmentation induction (D7, D14, D30), we analyzed enriched genes by population, cross-referenced the literature and annotated each cluster with a primary (group) and secondary (cluster) category. This process revealed a mixture of intermediate cell states resembling those described in the context of rostral embryo patterning and eye development (Bosze et al., 2020; Sarkar et al., 2020).

We named seven primary categories corresponding to key structures of the anterior ectoderm, such as stem cell-derived Lateral Neural Fold-like (LatNeEp), Pre-Placodal Epithelium-like (PrePlac), Cranial Neural Crest-like (CrNeCr), and Mesenchymal (MesCh), and lineages directly involved in eye morphogenesis, such as Retinal Progenitor (RetProg), Early RPE (EarlyRPE), and Intermediate RPE (MidRPE) (**Figure 2A**). Some secondary clusters matched remarkably well with specific neural tube regions, expressing a combination of enriched markers for eye field (*RAX, SIX6, LHX2*), telencephalic neural fold (*DLX5, DLX6*), lens placodes (*FOXE3, PAX6, ALDH1A1*), cranial neural crest (FOXC2, *VGLL2, PITX1*), inner ear placodes (*OTOGL, VGLL2, CYP26C1*), the anterior neural ridge organizer (*FGF8, SP8*, and *FOXG1*), and mesenchyme (*GABRP, HAND1, CO-L1A1*) (Cajal et al., 2012; Chen et al., 2017; Cohen-Salmon et al., 1997; Crespo-Enriquez et al., 2012; Gitton et al., 2011; Kasberg et al., 2013; Kumamoto and Hanashima, 2017; Seo et al., 2017; Soldatov et al., 2019; Tahayato et al., 2003) (**Figures 2B, S2A, and S2B; Table S1**).

**Figure 2.**
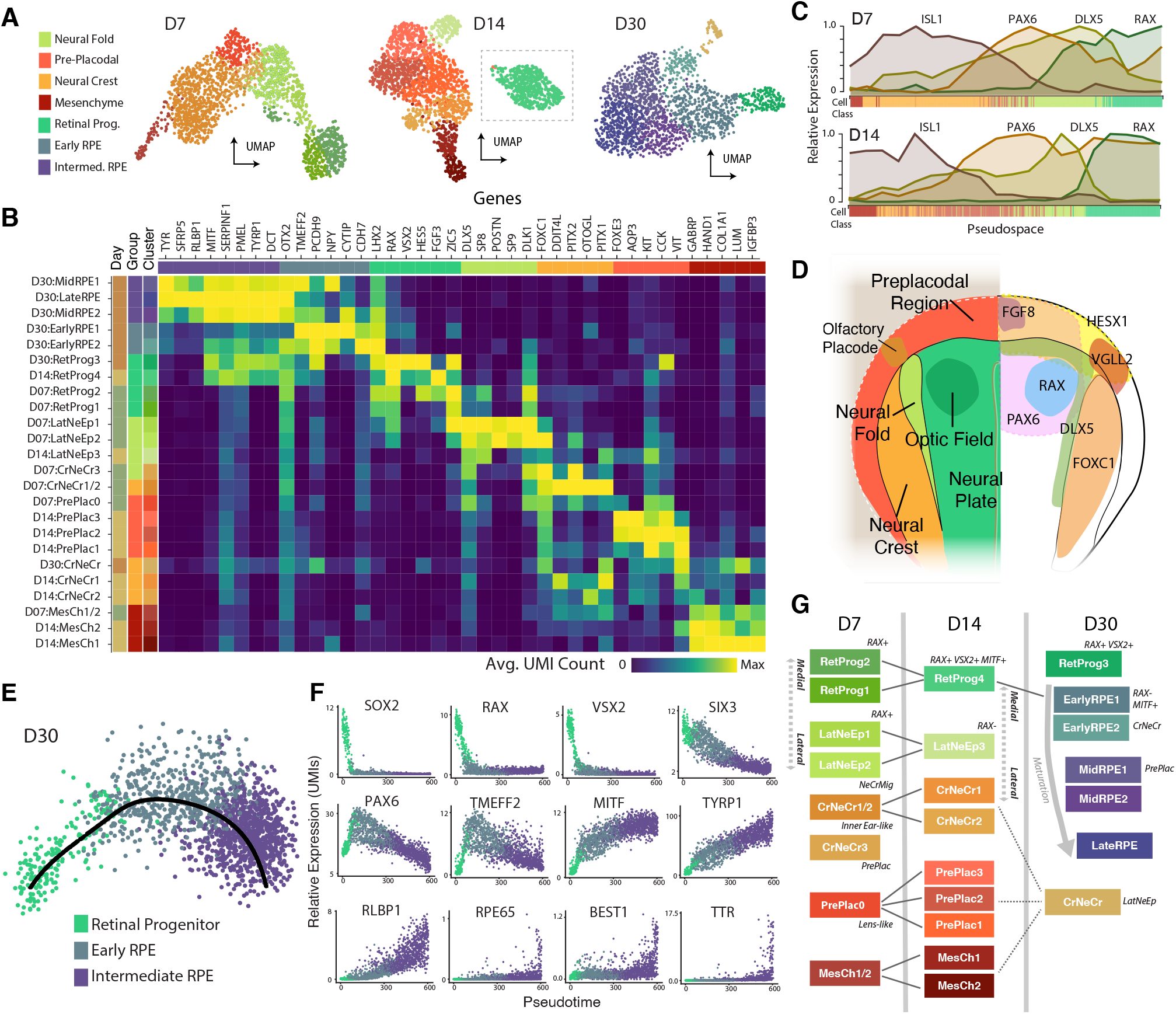
Evaluation of the diverse neuroepithelial cell type derivatives in early hESC-RPE differentiation. (A) UMAP representation of scRNA-seq at D7 (1,811 cells), D14 (1,872 cells), and D30 (1,852 cells). Clusters were grouped into seven primary categories: Lateral Neural Fold-like (LatNeEp), Pre-Pla-codal-like (PrePlac), Cranial Neural Crest-like (CrNeCr), Mesenchymal cells (MesCh), Retinal Progenitor (RetProg), Early RPE (EarlyRPE), and Intermediate RPE (MidRPE). (B) Enriched gene expression heatmap for hESC-RPE cell types. Heatmap clusters are annotated by differentiation day, primary group and secondary cluster. (C) Plots showing the relative gene expression of neural tube patterning markers *ISL1, PAX6, DLX5*, and *RAX* in D7 (top) and D14 (bottom) cells across the principal variation axis (pseudospace). Pseudospace axis was found by fitting a principal curve. The colored bar on the x-axis indicates the group assignment of single cells. (D) Schematic illustration summarizing the territories emerging in the early rostral embryo (left) and the gene expression patter of key genes (right). (E) Pseudotime trajectory of D30 RPE cells. (F) Progenitor (*SOX2, RAX, VSX2*, and *SIX3, PAX6*), early RPE (*TMEFF2, MITF*, and *TYRP1*) and late RPE (*RLBP1, RPE65, BEST1*, and *TTR*) gene expression along a pseudotime trajectory. (G) Schematic of the proposed relationship among the various secondary clusters during pigmentation induction. Edges indicate putative relationships between cell types identified at different time points. See also Figure S2 and Table S2.

The observation of *in vitro* MesCh is interesting because periocular mesenchyme expresses inductive signals *in vivo* that promote RPE fate, a role carried out by Activin A in the present protocol (Bosze et al., 2020). Eye field-like cells (RetProg) detected across days possessed distinct gene expression programs, suggesting varying degrees of progression towards RPE. D7 progenitors expressed Wnt antagonists *HESX1, FEZF1*, and *FRZB*, crucial for anterior fate assignment (Andoniadou et al., 2007; Peng and Westerfield, 2006) (**Figure S2C**). RetProg clusters expressed a repertoire of known markers, including *OTX2* and *LHX2*, which are jointly necessary for activation of the MITF transcription factor. These two genes were co-expressed in D14:RetProg4 alongside the *VSX2* neural retinal progenitor marker and MITF-activated genes *PMEL, SERPINF1, TYRP1* and *DCT*. D30:RetProg3 did not express MITF or downstream genes, suggesting these cells were less mature than D14:RetProg4. Consistent with their classification as progenitors, those cells displayed a stark cell proliferation signature (S and G2/M phases) and cell-cycle related RNA velocity (**Figures S2D and S2E**).

We next performed canonical correlation analysis (CCA) to integrate D7 and D14 cells on a shared feature space and factor out subtle time-dependent differences (**Figure S2F**). The joint representation captured a “pseudospatial” axis of variation, with cells transitioning along a mediolateral molecular profile (**Figures 2C, 2D, and S2G**). Over time, we observed an increase in cells assigned as lens-like pre-placodal (PrePlac; 15.19% to 48.04% of cells) and a decrease in both inner ear-like cranial neural crest (CrNeCr; 39.48% to 12.92%) and lateral neuroepithelial (LatNeEp; 21.48% to 6.41%) cells. The fraction of retinal cells was relatively unchanged (RetProg; 19.38% to 24.96%) (**Figure S2G**). These analyses revealed that expression heterogeneity at both stages recapitulates the molecular profile of rostral embryonic territories patterned to specify into sensory organs, such as lens, olfactory, and otic placodes (Begbie, 2013).

Conversely, a pseudotemporal trajectory of retinal maturation largely characterized D30 cells (see Methods; **Figure 2E**). Expression along a pseudotime confirmed a loss of progenitor status (*SOX2, RAX, VSX2, SIX3*), followed by an increase in RPE differentiation (*PAX6, TMEFF2, MITF, TYRP1*) and, later, of advanced RPE characteristics (*RLBP1, RPE65, BEST1, TTR*) (**Figure 2F**). Transcription factor network analysis with SCENIC further confirmed the activity of regulons involving factors *SOX2, RAX, VSX2, OTX2*, and *MITF* with anticipated gene targets (see Methods; **Figure S2H**). In fact, MITF cooperates with OTX2 to transactivate RPE pigmentation genes and downregulates progenitor genes (Martínez-Morales et al., 2003; Yun et al., 2009).

Our observations indicate that a sequential stepwise differentiation model is inadequate to explain the observed cell population dynamics; instead, the data suggests a “divergence-convergence” model with an initial expansion of cellular diversity, later dampened to favor the promotion of the RPE differentiation program (**Figure 2G**; see Discussion).

### *In vitro* hESC-RPE differentiation and eye development exhibit similarities in cellular composition and molecular profile

We next reasoned that embryonic references could validate our model and evaluate how faithfully *in vitro* phenotypes match their *in vivo* counterparts. Thus, we performed scR-NA-seq on an embryonic optic vesicle at 5 weeks post conception (W5; 2,637 cells) and two eyes at 7.5 weeks post conception (W7.5; 2,742 cells). Optic vesicle at W5 contained retinal epithelium more clearly differentiated into embryonic RPE (henceforth eRPE, to distinguish from the hESC-derived RPE clusters), neural retinal (eNR), and optic stalk (eOS) sub-populations than in the RPE-focused progenitors detected *in vitro* (**Figure 3A, cf. Figure 2B; Table S2**). In addition to retinal progenitor cells (eRetProg), we found two clusters (eCrNeCr, eMesCh) that co-expressed neural crest markers (Soldatov et al., 2019) (**Figures 3B and 3C**, right). The W7.5 eye captured a more diverse representation of cell types surrounding the eye, including proliferating progenitors, RPE, lens, intermediate retinal ganglion cells, and neural crest-derived mesenchyme (**Figures 3D, 3E, S3A, and S3B**). Progenitor markers highly expressed in W5 eRPE were more exclusive to neural (eNePr) and lens (eLensPr) progenitors by W7.5. Early differentiation genes were detected in eRPE, and a transition towards mature RPE was apparent in the embedding despite an absence of mature RPE markers *RPE65, BEST1*, and *TTR* (**Figures S3B and S3C**).

**Figure 3.**
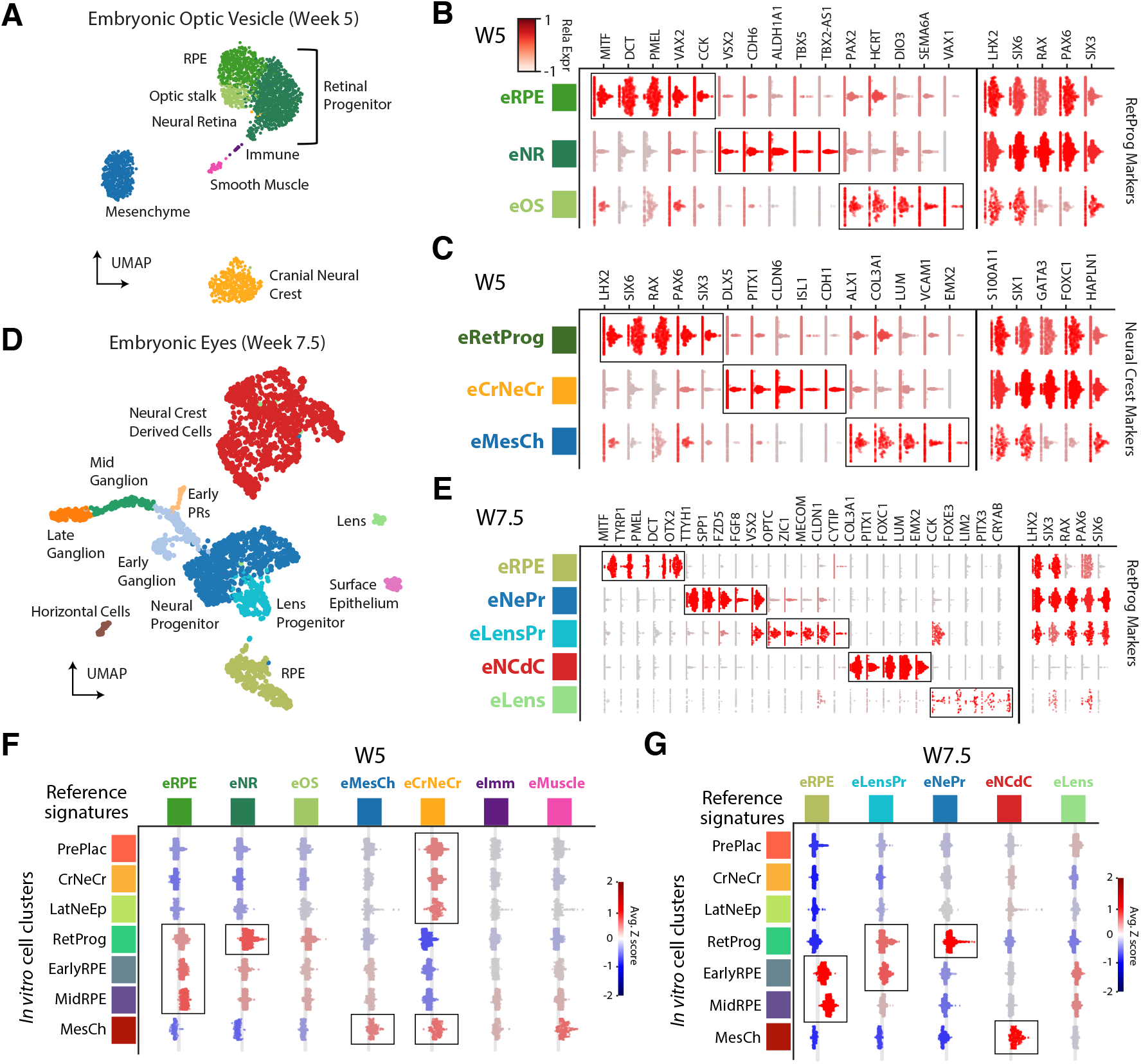
Comparative analysis of RPE induction between hESC-RPE and the human embryonic eye. (A) UMAP representation of a human embryonic optic vesicle (2,637 cells) dissected at 5 weeks (Carnegie Stage 13). Cluster identities include: optic cell types derived from retinal progenitors (eRetProg), such as retinal pigment epithelium (eRPE), neural retina (eNR), and optic stalk (eOS), in addition to periocular mesenchyme (eMesCh), cranial neural crest (eCrNeCr), immune (eImm), and smooth muscle (eMuscle). (B) Left: violin plots of enriched genes in eRPE, eNR, and eOS clusters at W5. Right: expression of broad retinal progenitor markers. (C) Left: violin plots of enriched genes in eRetProg, eCrNeCr, and eMesCh clusters at W5. Right: neural crest marker expression. (D) UMAP representation of two human embryonic eyes (5,274 cells) at 7.5 weeks. Cluster identities include: RPE (eRPE), neural progenitor (eNePr), lens progenitors (eLensPr), neural crest derived cells (eNCdC), and lens (eLens). (E) Left: violin plots of enriched genes in the identified W7.5 clusters. Right: retinal progenitor marker expression from (B) in eRPE, eNePr, eLensPr, eLens and eNCdC at W7.5. (F-G) Violin plots showing the distribution of gene expression signatures scores computed on D7, D14, and D30 hESC-RPE single cells and aggregated by the discovered clusters (rows). The signatures (columns) for to each of the embryonic cell types include the top 30 enriched genes for that cluster. See also Figure S3 and Table S3.

To evaluate the resemblance of hESC-RPE tissues to embryonic references, signatures were extracted from *in vivo* cell types and a score was calculated on *in vitro* cell groups at D7, D14, and D30 (see Methods; **Figures 3F and 3G**). The W5 eNR score was strongest for RetProg, and eRPE signatures from both W5 and W7.5 were associated with hESC-derived EarlyRPE and MidRPE. RetProg shared little resemblance to eRPE at W7.5 but was similar to eNePr and eLensPr. Overall, the molecular profiles of early hESC-RPE populations mirrored those present during development, but with a bias towards RPE fate over other retinal cell types.

### Cell surface marker NCAM1 defines retinal progenitor cells at D30 of hESC-RPE differentiation

Sorting out progenitor populations at intermediate stages could be both a route to faster RPE differentiation protocols and also a strategy to obtain a cellular source from which to derive other retinal lineages. Following the observation of a retinal progenitor cell population in D30 hESC-RPE cultures, we reasoned that reliable retinal progenitor markers would be inversely correlated to genes characterizing more mature cells, such as RPE, as well as to progenitors for other neuroepithelium tissues. We therefore computed a Pearson’s correlation coefficient between highly expressed genes at D30 and RPE or neural tube markers (see Methods; **Figures 4A and 4B; Table S3**). Genes with strong anticorrelation to both signatures encoded functionally diverse gene products, including cytoplasmic proteins, transcription factors, secreted molecules, and membrane proteins (**Figure 4C**). Transcription factors involved in early retina development (*CRABP1, RAX, ZIC2, SIX6*) ranked among the top genes, along with genes implicated in neural tube (*CPAMD8, PK-DCC, NR2F1*) and lens development (*MARCKS, DACH1, MAB21L1*) (Imuta et al., 2009; Yamada et al., 2003; Zhou et al., 2010).

**Figure 4.**
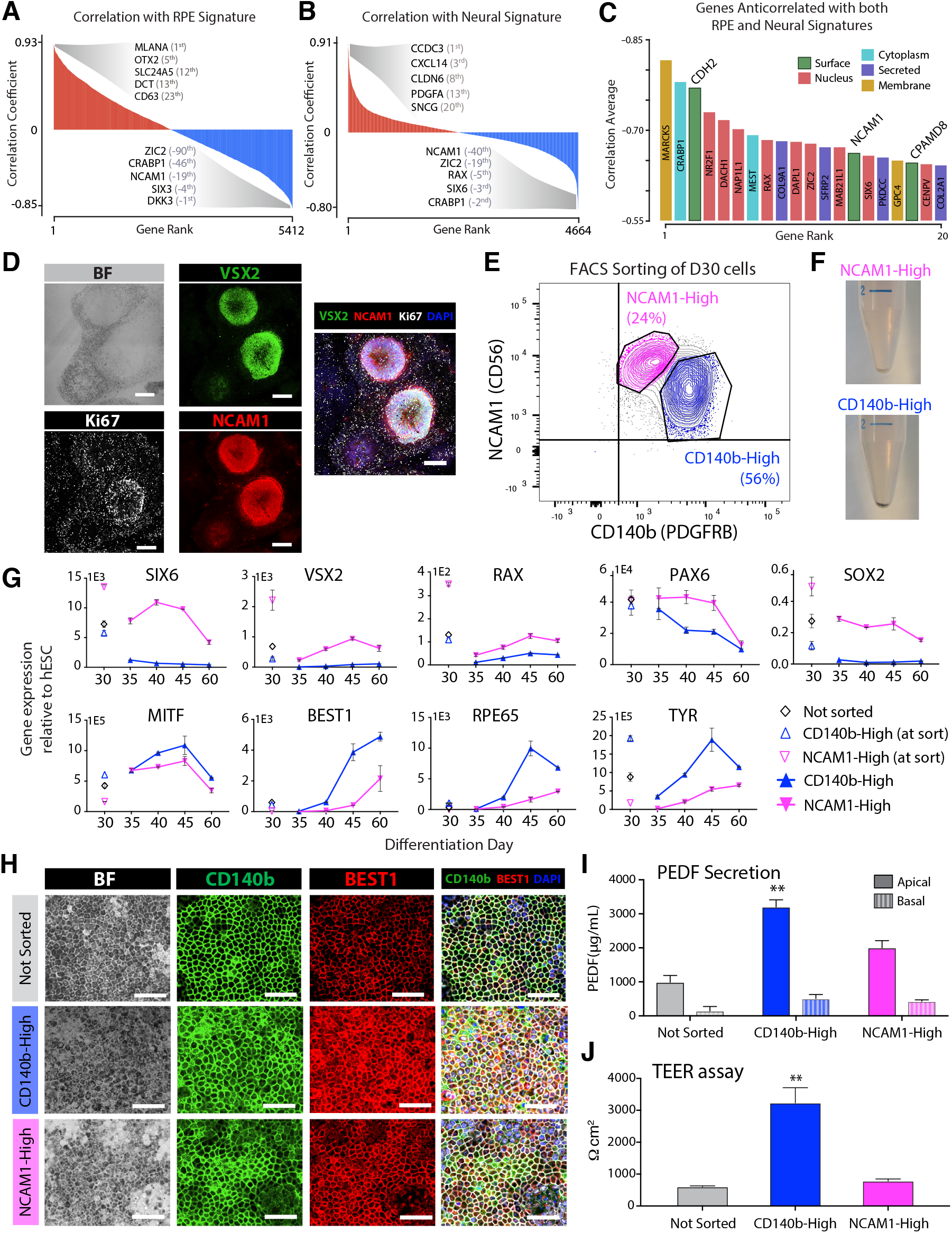
Characterization of NCAM1 sorted D30 hESC-RPE population. (A-B) Genes ranked by their correlation with two signature scores: an RPE signature (A) and a neural signature (B). Signature scores were computed using the top enriched genes in the RPE cluster and of the retinal progenitor and neural clusters. (C) Bar graph showing the top 20 genes ranked by the average (negative) correlation of their expression with the RPE and neural signatures (cf. A,B). Genes are colored by the annotated protein cellular localization. (D) Brightfield and immunofluorescence stainings of D30 showing co-expression of VSX2, NCAM1 and Ki67. Scale bars: 200μm. (E) Representative FACS plot of the NCAM1-CD140b sorting used to seprated CD140b-High and NCAM1-High populations at hESC-RPE D30. (F) Post-sort pellets of CD140b-High cells and NCAM1-High. (G) RT-qPCR of retinal progenitor (*SIX6, VSX2, RAX, PAX6, SOX2*) and RPE (*MITF, BEST1, RPE65, TYR*) marker genes in unsorted, CD140b-High and NCAM1-High populations at the moment of sort and at post-sort D30, 35, 40, 45, and 60. (H) Brightfield and immunofluorescence stainings of unsorted, CD140b-High and NCAM1-High populations 30 days after sorting (D60) showing co-expression of CD140b and BEST1 markers. Scale bars: 100μm. (I-J) PEDF secretion (I) and TEER measurements (J) of the unsorted, CD140b-High and CD56-High populations at D60 (30 days post-sorting). (G, I-J) Bars represent mean +/-SEM from three independent experiments. **p < 0.0001 (PEDF Apical, TEER) compared with the Not sorted and NCAM1-High conditions. See also Figure S4 and Table S4.

Interestingly, cell surface markers *CDH2, NCAM1*, and *CPAMD8* were among the strongest progenitor markers at D30. We were particularly intrigued by NCAM1 (surface antigen CD56), which was previously identified in a screen for eye field progenitor markers and whose expression was shown to be anticorrelated with pigmentation (Plaza Reyes et al., 2020a). In fact, NCAM1 staining areas coincided with rosette structures lacking pigmentation, and both scRNA-seq and protein staining revealed that *NCAM1* was co-expressed with progenitor (VSX2 and RAX) and proliferative (Ki67) genes at D30 (**Figures 4D, S4A, S4B, and S4C**).

This evidence suggested that NCAM1 may be a valuable progenitor-cell surface marker in differentiating RPE cultures. We therefore devised a sorting strategy to isolate this population using NCAM1 and CD140b (PDGFRB), an RPE cell marker (Plaza Reyes et al., 2020a). By combining the two markers, we separated cells into either a putative retinal progenitor stage (24% cells, CD140b^low^NCAM1^high^, henceforth NCAM1-High) or a more mature RPE stage (56% cells, CD140b^high^NCAM1^low^, henceforth CD140b-High) (**Figure 4E**). Consistently, a pigmented pellet was evident in the CD140b-High population, whereas the NCAM-1-High pellet lacked pigmentation, suggesting different intrinsic potentials and maturation statuses (**Figure 4F**). We then sorted D30 NCAM-1-High and CD140b-High cells and continued differentiation for the remaining 30 days. Morphological evaluation showed that CD140b-High cells generated a homogeneous hESC-RPE monolayer already at D45, while NCAM1-High cells instead yielded a defined RPE population only at D60 (**Figure S4D**).

We next assessed retinal progenitor (*SIX6, VSX2, RAX, PAX6, SOX2*) and RPE (*MITF, BEST1, RPE65, TYR*) markers by RT-qPCR to understand the expression dynamics corresponding to our morphological observations. The measurements confirmed that NCAM1-High cells expressed higher levels of progenitor genes than CD140b-High cells at the time of sorting. Progenitor genes continued to be present after sorting in NCAM-High cultures (declining over time under RPE differentiation conditions), whereas in CD140b-High cultures the interrogated genes were close to absent throughout the protocol. Conversely, CD140b-High cells upregulated mature RPE markers earlier and more rapidly than NCAM1-High cells (**Figure 4G**).

The percentage of cells positive for the cell cycle marker Ki67 was higher in the NCAM1-High population, indicative of their immature and proliferative state, and eventually declined upon reaching an RPE phenotype (**Figure S4E**). In fact, VSX2 protein was more highly abundant in NCAM1-High cells that also co-expressed Ki67 for a longer time under RPE conditions than in CD140b-High cells (**Figure S4F**). Establishment of an RPE phenotype at D60 by both NCAM1-High and CD140b-High populations was confirmed by the appearance of a cobblestone morphology and pigmented cultures in addition to co-expression of CD140b and BEST1 proteins (**Figure 4H**).

To evaluate the functional relevance of those changes and compare the degree of differentiation between the two populations, we assessed pigment epithelium-derived factor (PEDF) secretion and transepithelial resistance (TEER) upon protocol completion (D60), finding that CD140b-High-derived cells secreted significantly higher apical levels of PEDF than the unsorted and NCAM1-High-derived cells, both at similar standard levels (1000-2000 ug/mL) (**Figures 4I**). TEER levels displayed by CD140b-High-derived cells were significantly superior to unsorted and NCAM1-High populations, whose levels were comparable to D60 hESC-RPE cells (400-800 Ω*cm2) (Plaza Reyes et al., 2016, 2020a) (**Figure 4J**). These results show that CD140b selects for more quickly maturing pigmented cells at D30, whereas NCAM1 denotes an immature population with progenitor potential that is eventually, under specific RPE-driving cues, also capable of generating functional RPE.

### NCAM1-High cells can differentiate into alternative retinal cell types

To evaluate the differentiation potential of NCAM1-High cells, the population was sorted at D30 and plated in neural retinal progenitor-promoting conditions for 40 additional days (Shao et al., 2017) (**Figure 5A**). Interestingly, NCAM1-High cells gave rise to a more heterogeneous culture, with a significant portion of cells displaying a distinct non-RPE cell body morphology (**Figure 5B**). Gene enrichment analysis of 980 single cells yielded a variety of molecularly-distinct populations, of which only 12% were RPE, confirming that NCAM1-High cells at D30 represent an uncommitted progenitor (**Figure 5C**).

**Figure 5.**
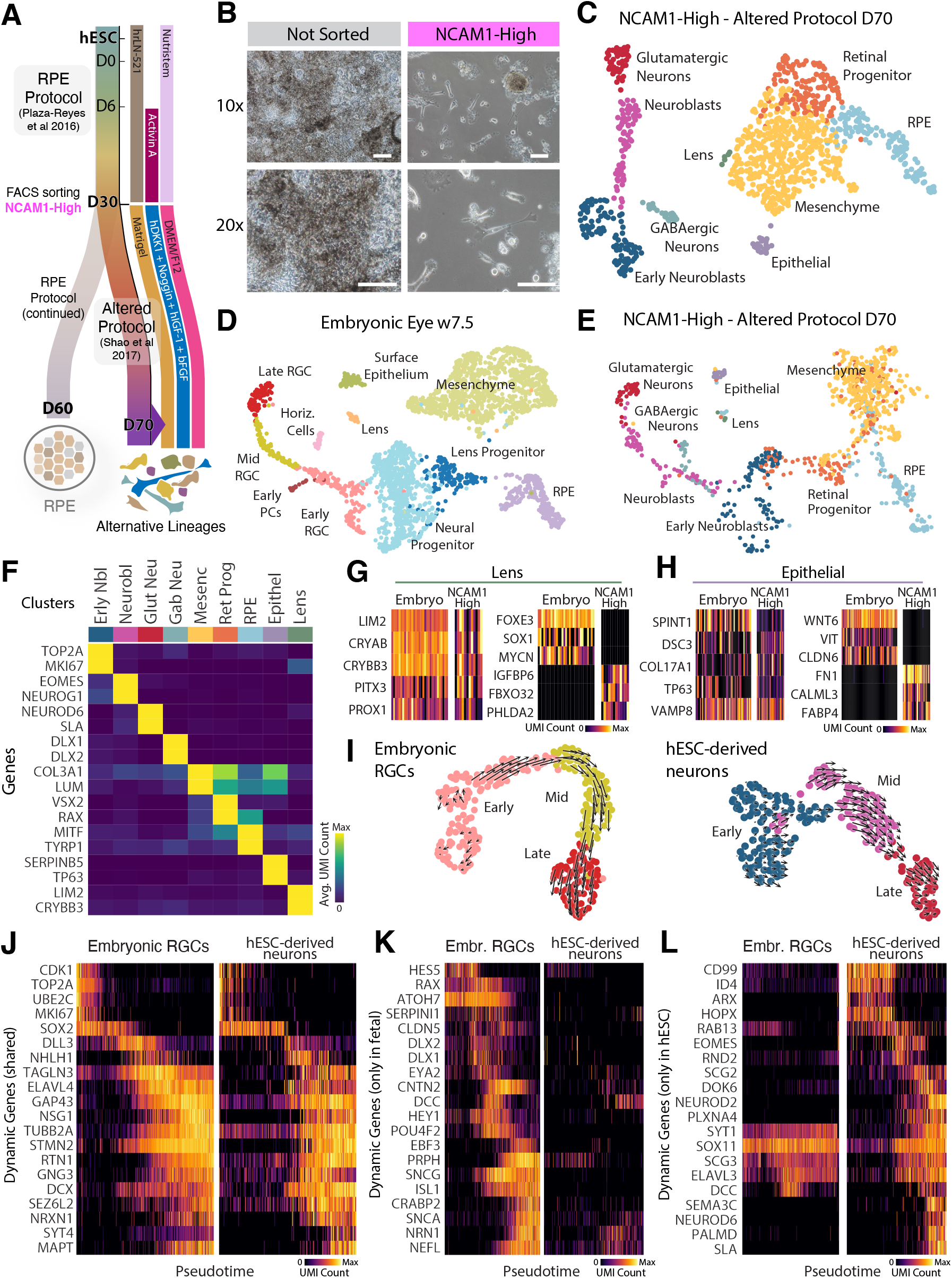
Altered differentiation of NCAM1-High sorted hESC-RPE D30. (A) Schematic of the neuronal differentiation protocol. hESC-RPE differentiation was performed up to D30; NCAM1-High cells were then sorted and replated on matrigel containing DMEM/F12, hDKK1, Noggin, hIGF-1, and bFGF until scRNA-seq at D70. (B) Brightfield images of unsorted and sorted NCAM1-High populations at D70 following the altered protocol. Scale bars: 100μm. (C) UMAP representation of NCAM1-High sorted hESC-RPE at D70 (980 cells) following the altered protocol, including the following cell types: early neuroblasts (Erly Nbl), neuroblasts (Neurobl), glutamatergic neurons (Glut Neu), GABAergic neurons (Gab Neu), mesenchymal cells (Mesenc), retinal progenitor (Ret Prog), RPE, epithelial (Epithel), and lens (Lens). (D-E) Integrated embedding of the scRNA-seq data from embryonic W7.5 eye (D) and NCAM1-High hESC-RPE cells subjected to the altered protocol (E). (F) Enriched gene expression for cell types identified in (C). (G-H) Gene expression heatmaps for rare Lens (G) and Epithelial (H) cells identified in the embryo and *in vitro*. Shared markers are shown on the left plots; differential expressed genes in the plots on the right. (I) Left: RNA velocity analysis of embryonic retinal ganglion cells, classified as Early, Mid, and Late. Right: RNA velocity of hESC-derived neurons, classified as Early (Erly Nbl, cf 5C), Mid (Neurobl, cf 5C), and Late (Glut Neu, cf 5C) (J-L) Gene expression analysis of embryonic and hESC-derived neuronal along their respective pseudotimes. RGC (Retinal Ganglion Cell), PC (Photoreceptor cell), Horiz. cells (Horizontal cells).

We then performed CCA integration with the W7.5 embryonic eyes to systematically compare NCAM1-High-derived cells to a developmental reference. The shared low dimensional space emphasized similarities between corresponding clusters, including RPE, progenitor, mesenchymal, lens, surface epithelial and neuronal populations (**Figures 5D, 5E, and 5F**). Non-RPE retinal cell types detected included a small lens population co-expressing *LIM2, CRYAB, PITX3*, and *PROX1*. Genes exclusive to embryonic lens (*FOXE3* and *SOX1*) are specific to promoting early lens development, suggesting that the NCAM1-High-derived lens cells are in a more mature state (Blixt et al., 2000; Nishiguchi et al., 1998) (**Figure 5G**). A shared epithelial population was also observed, co-expressing markers characteristic of surface epithelium and keratinocytes, which can be found in the cornea (**Figure 5H**).

Moreover, there was an overlap between W7.5 retinal ganglion neurons and the NCAM1-High-derived neurons (**Figures 5D and 5E, cf. Figure S3**). To compare gene expression dynamics of these cells, RNA velocity was computed on each neuronal population, revealing progression towards a more mature state (**Figure 5I**). Pseudotemporal gene expression confirmed a common profile of expression waves, with gradual downregulation of proliferation markers (*TOP2A, MKI67*) followed by upregulation of a neuronal differentiation program (*TAGLN3, STMN2, TUBB2A, DCX, NRXN1*) (see Methods; **Figure 5J**). However, crucial markers of retinal ganglion development, such as transcription factor *ATOH7* and its downstream targets *POU4F2* and *ISL1*, were only expressed in the embryonic cells (Gao et al., 2014) (**Figure 5K**). Other neuronal markers (*EOMES, NEUROD2, NEUROD6, SLA*), were unique to NCAM1-High-derived neurons, implying that NCAM1-High-derived cells are another type of telencephalic neuron (**Figure 5L**). NCAM1-High cells are thus either a mixed pool of retinal and neuroepithelial progenitors capable of forming both cell types and other related retinal lineages, or cells with the capacity to retain the potential of all these lineages (**Figure S4A**; see Discussion).

### Late hESC-RPE differentiation is characterized by the selection and maturation of RPE populations

We next analyzed scRNA-seq at three subsequent time points after D30 replating (D38, D45, D60). Unlike the initial stages, most annotated clusters consisted of RPE. Indeed, the proportion of RPE cells increased from 82.1% at D38 to 98.7% at D60, confirmed by expression of early differentiation markers *MITF* and *TYRP1* as well as mature pigmentation and visual cycle markers *RLBP1, RPE65, BEST1*, and *TTR*, which were detected only in a small proportion of cells at early time points. We also annotated small fractions of non-retinal types, including smooth muscle mesenchyme (*VTCN1, HAND1, WNT6*) and myogenic contaminants (*ACTA1, MYOD1, MYOZ2*) (**Figures 6A, 6B and S5A**). Interestingly, the D38 time point after replating displayed an increased heterogeneity and was on average less differentiated than D30 (**Figure 1**). Surprisingly, D38 contained a small cluster expressing stem cell markers (*POU5F1, LI-N28A, SALL4*) alongside neural crest (PlrNeCr, 2.4% cells) genes (*TJP3, FZD2, FOXI3*) notably absent in undifferentiated hESCs (**Figure S5B**). Furthermore, we observed from D30 onwards a population of RPE co-expressing MITF and markers associated with the epithelial-to-mesenchymal (EMT) transition process, particularly *ACTA2*. Recent studies have suggested that TGFβ signals used in RPE differentiation protocols can inadvertently induce EMT, whose markers are co-expressed with *MITF* (Boles et al., 2020; Jung et al., 2020; Salero et al., 2012). Despite the absence of Activin A in culture from D30 and onwards, the dissociation of RPE cells at D30 nonetheless induced a mesenchymal-like morphology of the RPE cells (**Figure 1B**). This finding led to characterization of maturing RPE (*MITF*+*ACTA2*-) and EMT-RPE (*MITF*+*ACTA2*+) (**Figure S5C**). The proportion of EMT-RPE increased during replating from D30 to D38, followed by a steady decrease to low levels (1.2% cells) at D60 (**Figures S5D and 6C**). These cells displayed a signature of some, but not all, RPE markers co-expressed with EMT markers. Moreover, the representation of RPE from later time points along a phenotype variation axis confirmed the presence of some shared EMT and RPE differentiation properties (**Figures S5E and 6D**). We concluded that replating might select against more differentiated cells, activate a temporary expansion of progenitors, or both. Nonetheless, in the final 30 days, we observed the persistence of RPE and loss of other cell types, with 98.7% cells at D60 of RPE fate and remaining 1.3% retinal progenitors (**Figures 6C and 6D**). In addition to the 1.2% of EMT-RPE, there was some maturation variability among the RPE clusters at D60, and in fact, pseudotime trajectory inference and RNA velocity analysis confirmed the movement of less mature populations in gene expression space towards the most mature RPE (**Figures 6E and 6F, cf. Figure 3F**). CCA integration of D60 cells with a second protocol replicate (931 cells) from prior study (Plaza Reyes et al., 2020a) confirmed the reproducibility of the differentiation procedure, final cell type proportions, and gene expression patterns (**Figures S5G and S5H**). Moreover, the analysis of the gene expression correlation matrix of all hESC-RPE differentiation clusters helped to further chart the relationships across time points, reemphasizing the global stem cell to hESC-RPE transformation during the differentiation protocol and highlighting similarities among clusters of comparable maturation states at different time points (**Figures S5A and S5I**).

**Figure 6.**
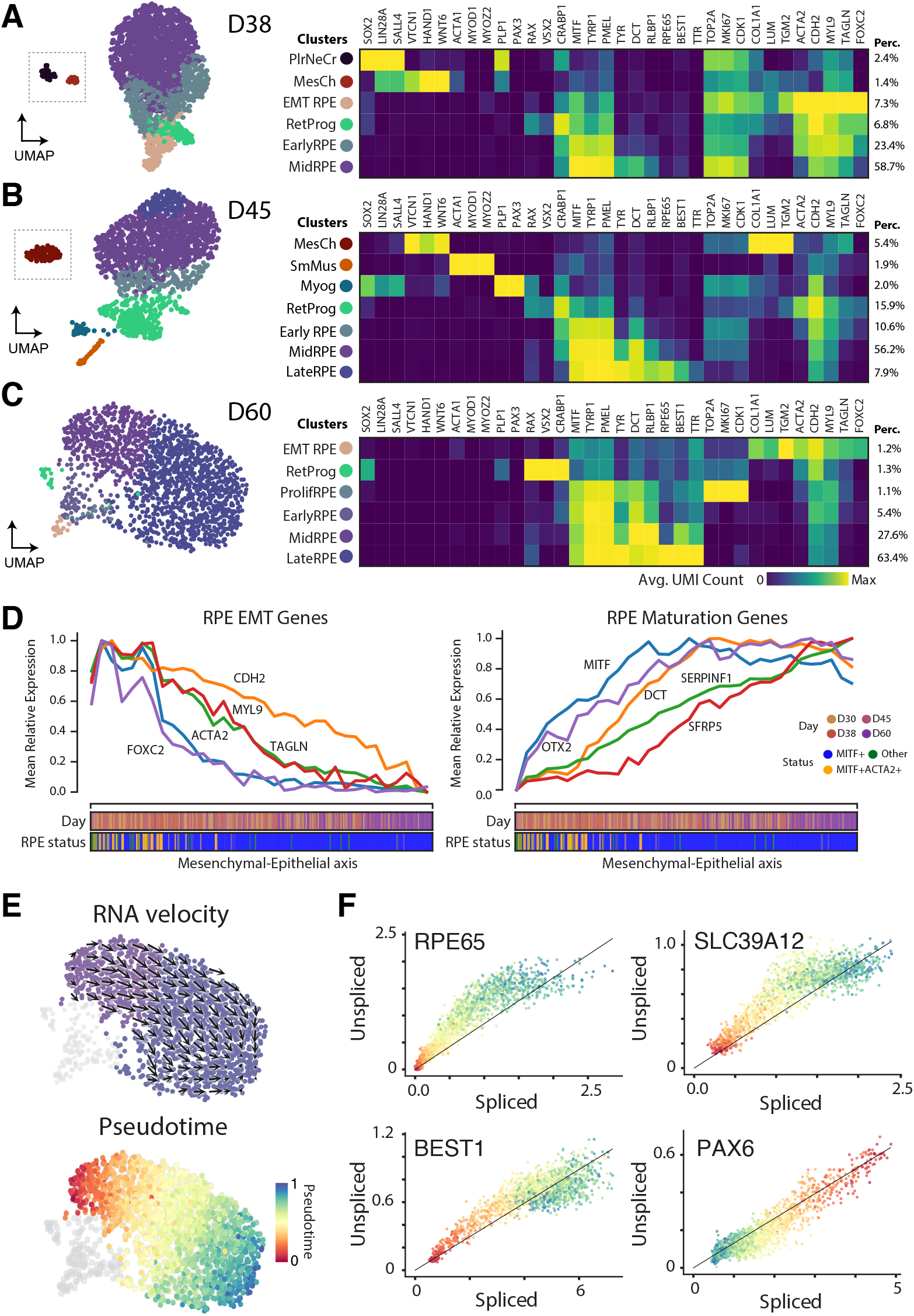
Late hESC-RPE differentiation profiling. (A-C) UMAP representations and marker gene expression heatmaps of 1,784 cells at hESC-RPE D38 (A), 1,707 cells at hESC-RPE D45 (B), and 1,749 cells at hESC-RPE D60 (C). Cell-type proportions are indicated on the right. (D) Line-plots showing average expression of RPE-EMT (left) and mature RPE (right) along a mesenchymal-epithelial axis of variation determined fitting a principal curve (See Methods). Colored bars on the x-axis indicate timepoint and RPE status of cells along the axis. (E) Top: RNA velocity analysis of Mid and Late RPE populations at hESC-RPE D60. Bottom: The same cells colored by their pseudotime assigment. (F) RNA-velocity phase portraits scatters. The concave shape of the scatters of RPE marker genes *RPE65, BEST1*, and *SLC39A12* is evidence of an ongoing upregualtion dynamics while the convex shape of the scatter of the progenitor marker *PAX6* of an ongoing downregulation. See also Figure S5.

### Subretinal transplantation of hESC-RPE facilitates a more advanced RPE state

Robust hESC-RPE differentiation is a required first step towards cellular therapies for retinal degeneration. However, derived RPE cells must integrate with neighboring tissues upon injection, retain mature attributes and avoid the resurgence of pluripotent properties to become an effective treatment modality. To evaluate these aspects, D60 hESC-RPE cells were transplanted in the subretinal space of two albino rabbits, a preclinical large-eyed animal model (Bartuma et al., 2015; Petrus-Reurer et al., 2017, 2018). Transcriptional analysis of adult rabbit (1,965 cells) and adult human retina (5,538 cells) showed a high degree of similarity (**Figure S6; Table S4**). Four weeks following transplantation of hESC-RPE, infrared and SD-OCT imaging showed a pigmented patch of human cells and a hyper-reflective RPE layer among the albino rabbit retinal layers (**Figure 7A**). Histology and immunofluorescence staining further demonstrated co-expression of pigmentation, human marker NuMA and the RPE marker BEST1, corroborating the successful integration of injected hESC-RPE cells in a polarized monolayer (**Figure 7B**). The contiguous injected retina of two rabbits was then processed for scRNA-seq, yielding 65 human hESC-derived cell profiles exhibiting strong expression of mature RPE markers. Crucially, markers of retinal progenitors, photoreceptors, pluripotent hESCs, and EMT-RPE were benchmarked against our references and found to be absent following transplantation (**Figures 7C**). These findings attest that integrated hESC-RPEs possess the unique transcriptional signature of mature RPE without signs of retinal progenitor or pluripotent properties.

**Figure 7.**
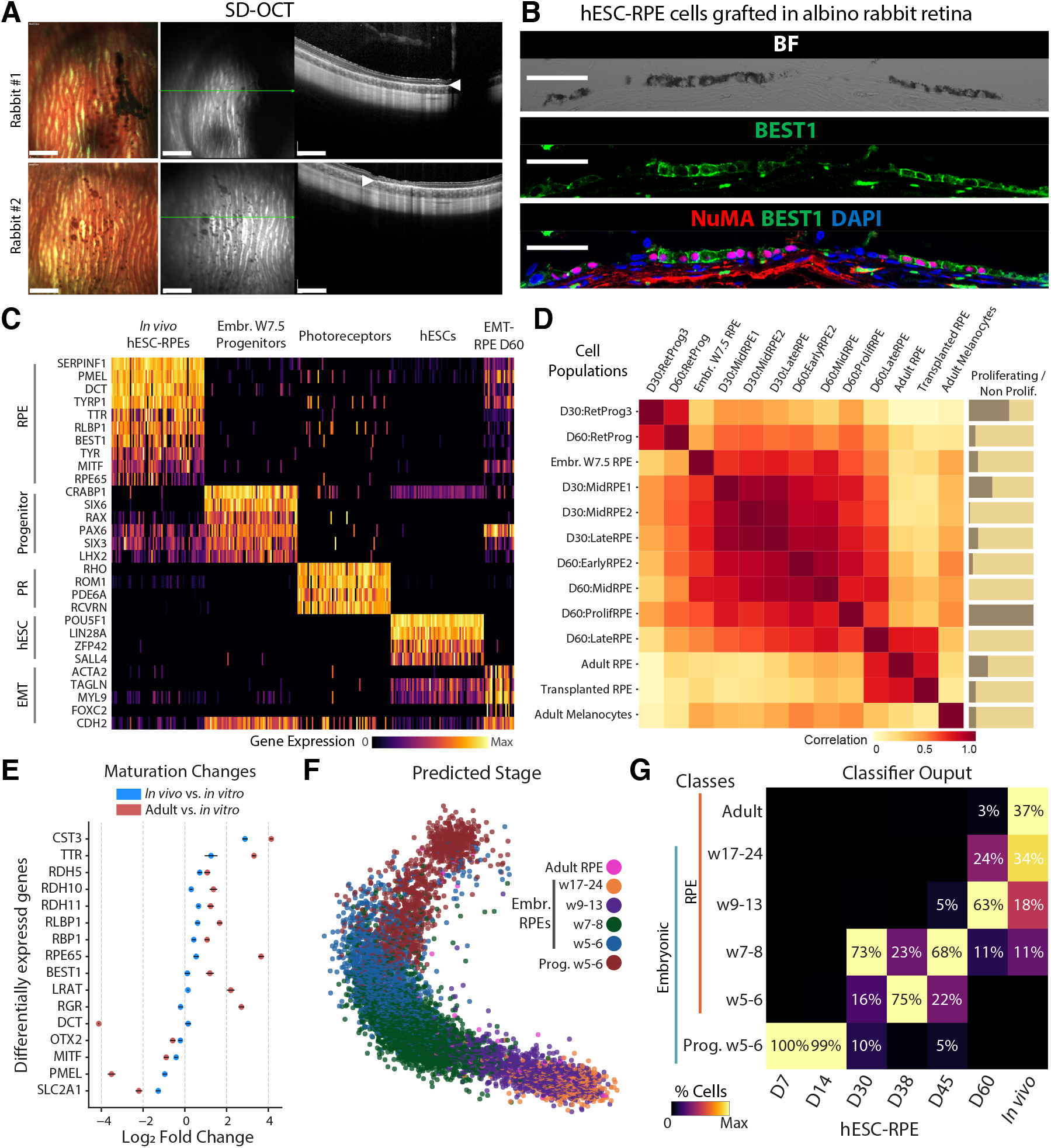
Phenotyping of hESC-RPE transplanted in the albino rabbit subretinal space. (A) Infrared and SD-OCT images of injected hESC-RPE D60 cells into the subretinal space of albino rabbits. Green lines indicate the SD-OCT scan plane. White arrows indicate the hyper-reflective RPE layer. Scale bars: 1mm. (B) Brightfield and immunofluorescent staining for NuMA and BEST1 30 days after hESC-RPE injection. Scale bars: 50μm. (C) Gene expression heatmap comparing 65 single hESC-RPE cells integrated in the rabbit subretinal space for 30 days to embryonic W7.5 retinal progenitors, adult photoreceptors, undifferentiated hESCs, and D60 EMT-RPE. (D) Pearson’s correlation matrix between the gene expression profiles of hESC-RPE D30 and D60, *in vivo* transplanted RPE, adult RPE and melanocytes, and embryonic RPE. (E) Dot-plot showing the fold change of crucial RPE markers between hESC-RPE D60 cells, transplanted RPE, and adult RPE. Bars represent mean +/-SEM from all cells at each time point. (F) Scatterplot of the UMAP embedding of *in vitro* hESC-RPEs from D7 to D60 (cf. Fig. 1C) colored by the classification results of na ordinal classifier fir on the embryonic developmental scRNA-seq data from Hu et al, 2019 (embryonic weeks 5-24). (G) Classification summary matrix showing the percent correspondence between predicted RPE developmental time point (embryonic weeks 5-24), adult RPEs, hESC-RPE timepoints (D07, D14, D30, D38, D45, D60) and transplanted RPE in the rabbit subretinal space (*In vivo*). See also Figures S6, S7, and Table S5.

We further compared *in vivo* and *in vitro* expression patterns by performing gene expression correlation analysis using both RPE and RetProg clusters at D30 and D60 as well as embryonic reference tissues (**cf. Figures 3, S3, S6**). Most RPE clusters at D30 and D60 were well correlated, yet only the most mature *in vitro* cluster (D60:LateRPE) and the *in vivo* transplanted RPE were highly similar to the adult RPE reference. Melanocytes, a distinct neuralcrest derived cell type with similar marker expression to RPE, were not as correlated with hESC-RPE. Interestingly, while D60:LateRPE retained similarities with other *in vitro* clusters, the transplanted RPE did not (**Figure 7D**). Differential expression analysis confirmed an expression pattern closer to adult RPE cells after *in vivo* implantation, particularly for visual cycle components such as *TTR, RPE65, RBP1, RLBP1*, and *RDH5* (**Figures 7E, S7A, and S7B**).

Lastly, to assess overall progression of hESC-RPE and assign cells to developmental stages, we built an ordinal classifier using 783 single-cell transcriptomes from human W5 to W24 (Hu et al., 2019) (**Figures S7A and S7B**). Our classifier operates with an underlying knowledge of the sequential relationship among the training data and was applied to place hESC-RPE cells on the temporal spectrum of RPE development (see Methods). We observed a gradual progression of maturity during *in vitro* differentiation corresponding to embryonic RPE development. Transplanted hESC-RPE were assigned to the adult RPE class more than any *in vitro* time point (**Figures 7F and 7G**). As a proof of principle, we applied the classifier to our human embryonic references at W5 and W7.5, confirming an appropriate assignment (**Figures S7C and S7D**). Importantly, grafted cells also exhibited similar levels of apoptotic and stress response genes to ungrafted cells (**Figures S7E and S7F**). Overall, despite the pluripotent state of the cell source and the initial diversity expansion observed, the hESC-RPE differentiation protocol ultimately yielded homogenous and mature RPE cells in a sequence similar to that of embryonic RPE development, and subretinal transplantation assisted with further maturation towards a more adult state.

## DISCUSSION

In the present study, by molecularly profiling a directed and defined hESC-RPE differentiation protocol established for clinical translation (Plaza Reyes et al., 2020a), we demonstrated that over 60 days, culture conditions successfully induced RPE lineage specification and maturation. We observed a sequence of gene expression waves consistent with embryological studies (Fuhrmann et al., 2014; Hu et al., 2019). However, at early stages we found a cell pool heterogeneity that was incompatible with the induction of a single lineage and instead was more consistent with an initial expansion of cellular diversity (**Figures 1, 2, and S6A**). Similar cell type heterogeneity expansion was observed in studies of endoderm and endothelial tissue derivation, but meta-analyses of several differetiation protocols will have to be performed to understand to which extent such initial diversity expansion is a widespread phenomenon (Cuomo et al., 2020; MacLean et al., 2018; McCracken et al., 2020).

In the analyzed derivation protocol, we mainly identified cell populations reminiscent of different rostral embryonic tissues. In addition to eye field progenitors, we found expression profiles resembling patterned regions surrounding the optic field: the pre-placodal epithelium, neural fold, and neural crest (**Figure 2B**). These intermediate types are induced in the embryo by the anterior neural ridge organizer, which promotes specification of the sensory placodes, including the lens ectoderm surrounding the optic cup (Begbie, 2013; Eagleson et al., 1995; Fuhrmann, 2010; Streit, 2007). We also observed cells expressing inner ear and olfactory-related genes, highlighting the closely-linked repertoire of transcription factors driving sensory tissue specification (Saint-Jeannet and Moody, 2014; Singh and Groves, 2016). These expression patterns and the RPE maturation timeline was further corroborated through comparisons to embryonic eye references (**Figures 3 and S3**). Taken together, characterization of our *in vitro* hESC-RPE differentiation process suggests a divergence-convergence model: a diversity expansion at early time points followed by selection of RPE lineage, driven by protocol conditions, and convergence onto a homogeneous and pure cellular product.

These findings hint at an intriguing self-organization process occurring in our 2D culture, despite the lack of spatially directed cues or 3D structure. An early spontaneous generation of cell fates related to the target cell type has been described in studies investigating the composition of intermediate stages with unbiased approaches such as scRNA-seq (Cuomo et al., 2020; La Manno et al., 2016). Ultimately, these observations could be contextualized and expanded upon with a comparison between 2D and 3D culture models, as some reports suggest that a highly spatially organized system is not necessarily the most molecularly patterned (Quadrato et al., 2017; Velasco et al., 2019).

Finding the mechanism of induction for the molecular patterning observed is a direction for further investigation. For example, it might be interesting to evaluate the potential involvement of neural crest-like and mesenchymal cells appearing during the protocol in the differentiation process, given their known endogenous roles *in vivo* (Fuhrmann et al., 2000; Kagiyama et al., 2005). There is a risk that the presence of many diverse rostral cell populations delays the maturation of a pure RPE target, a hypothesis consistent with the loss of maturity observed after replating and Activin A removal, especially from D30 to D38. Indeed, small smooth muscle and myogenic populations with co-expression of certain pluripotency markers temporarily emerged at D38, before culture conditions seemingly selected against such populations and facilitated a complete convergence on mature RPE by D60 (**Figures 6 and S5**). The emergence of cells with pluripotent signatures at D38 was surprising considering that there were no such cells detected at the earlier stages of differentiation. However, the finding emphasizes the importance of having robust methods to ensure that the final cell product does not contain lingering pluripotent cells that could lead to tumor formation.

Nonetheless, efficient sorting procedures for purification of the intermediate D30 population, prior to the emergence of such contaminants, may facilitate the design of a faster and purer differentiation protocol. We showed that removing the NCAM1-High progenitor pool at D30 yields a more mature RPE population in less time (**Figures 4 and S4**). Interestingly, we also demonstrated that NCAM1-High cells are not RPE-fate restricted and, upon altered culture conditions, give rise to different cell type lineages, including anterior neurons, mesenchyme, and lens epithelium (**Figure 5**). This potency is particularly relevant, as the identification and isolation of less mature progenitors with an increased plasticity is of importance to efforts aimed at replacement of other retinal cell types affected by advanced AMD (Bhatia et al., 2010; Marquardt et al., 2001). Thus, further evaluation of the NCAM1-High potency as a response to different and more specific culture conditions constitutes a promising avenue for future investigation.

The behavior of grafted cells *in vivo* is a topic discussed extensively by the community, with maintenance of the proliferative potential and dedifferentiation generally considered the two processes of major concern (Wang et al., 2020; Zarbin et al., 2019). Our analysis identified neither specific signs of dedifferentiation nor the presence of a non-RPE molecular profile. Instead, we detected a distinct shift in the RPE maturation towards a more adult and functional phenotype (**Figure 7**). The induction mechanism of this maturation remains unclear, however, the increased expression of visual cycle genes suggests that donor cells support neighboring photoreceptors functionally. Furthermore, after 30 days from the injection in the rabbit we did not observe any signs of stress-response in the grafted or the resident retinal cell types, suggesting that in a human niche with controlled alloimmune reaction, an increased survival and integration rate could be achieved (**Figures S7E and S7F**).

These observations encourage future attempts of injecting hESC-derived RPE at earlier stages of maturation, relying on the retinal microenvironment to induce further differentiation. In this context, extended characterization of the potency and differentiation capacity of different cell subpopulations becomes crucial, as they could constitute the candidate cellular subtype for further transplantation experiments. For example, exploring the capacity of the NCAM1-High progenitors in an *in vivo* model of retinal degeneration to specifically characterize their reparative potential.

Overall, our findings provide a high-resolution perspective on human pluripotent stem cell differentiation and a comprehensive and necessary detailed analysis of a stem cell-based product intended for successful and safe human therapeutic strategies. Ultimately, this study will guide future efforts focused on the differentiation of retinal cells, a deeper understanding of mechanisms of retinal disease, and applications in regenerative medicine.

## ACKNOWLEDGEMENTS

We thank Ernest Arenas, Alvaro Plaza Reyes, Pierre Fabre, Igor Adameyko, Pierre Gonczy, Felix Naef and Bart Deplancke for helpful feedback on the manuscript. The work performed in the Lanner laboratory was supported by grants from the Swedish Research Council, Ragnar Söderberg Foundation, Ming Wai Lau Center for Reparative Medicine, Center for Innovative Medicine, Wallenberg Academy Fellow, Strategic Research Area (SRA) Stem Cells and Regenerative Medicine. The Kvanta laboratory was supported by Stockholm County Council (ALF project), Karolinska Institute, Crown Princess Margaretas Foundation for the Visually Impaired, Strategic Research Area (SRA) Stem Cells and Regenerative Medicine, ARMEC Lindeberg Foundation, the Ulla och Ingemar Dahlberg Foundation, and King Gustav V and Queen Victoria Foundation and Cronqvist Foundation. G.L.M and A.R.L. were supported by the Swiss National Science foundation grants CRSK-3_190495 and PZ00P3_193445.

This study was performed at the Live Cell Imaging unit/ Nikon Center of Excellence, BioNut, KI, supported by Knut and Alice Wallenberg Foundation, Swedish Research Council, Centre for Innovative Medicine and the Jonasson donation. Flow cytometry analysis and cell sorting were performed at the MedH Flow Cytometry core facility, and prenatal human tissue was acquired through the Developmental Tissue bank core facility, both supported by KI/SLL. Sequencing was performed at ESCG Infrastructure in Stockholm at Science for Life Laboratory (funded by the Knut and Alice Wallenberg Foundation and the Swedish Research Council) with assistance from SNIC/Uppsala Multidisciplinary Center for Advanced Computational Science with massively parallel sequencing and access to the UPPMAX computational infrastructure.

## AUTHOR CONTRIBUTIONS

S.P.-R., A.R.L., F.L., G.L.M. conceived the study and planned experiments; F.L., G.L.M. supervised the work. S.P.-R., L. B.-V., M.W., H.A., E.S., and A.B. performed experiments; H.A., E.S., A.Kv., and A.Kr. contributed the human tissue datasets. I.D. and B.P. helped with the cell sorting. H.B., M. A. and A.Kv. contributed to the animal work; A.R.L. and G.L.M. performed the single cell RNA sequencing analyses; A.R.L., S.P.-R., A.Kv., G.L.M. and F.L. analyzed data. S.P.-R., A.R.L., G.L.M. and F.L. wrote the manuscript with input from all the coauthors.

---

**Figure S1.**
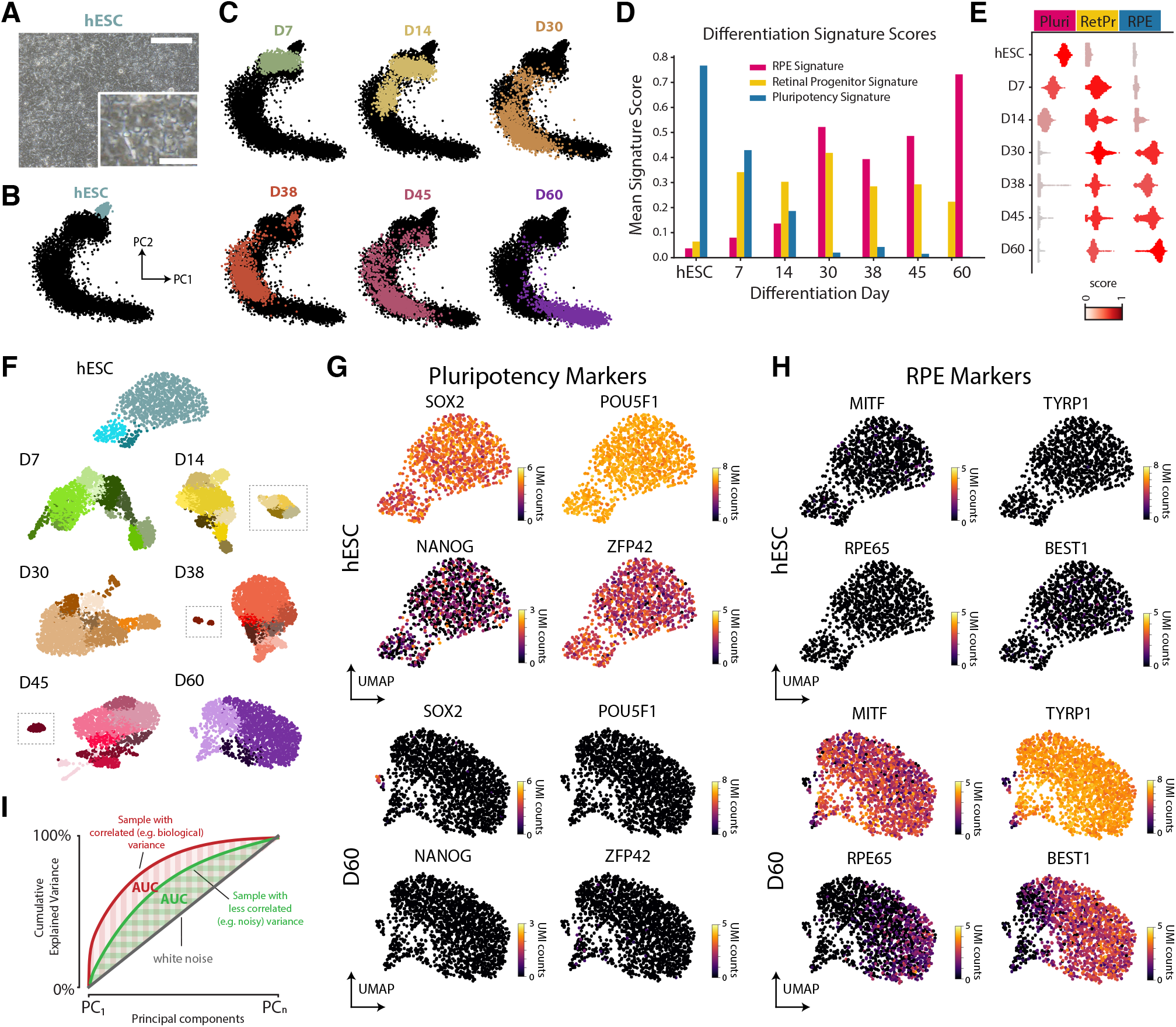
Cellular heterogeneity analysis of hESC-RPE differentiation, Related to Figure 1. (A) Brightfield image of undifferentiated hESCs. Scale bars: 100μm; inset 20μm. (B) Principal component representation of hESC. (C) Principal component representation of in vitro hESC-RPE time points coloured by day (see Figure 1). (D) Bar plot reporting average signature scores at each time point. (E) Violin plot indicating the distribution of the signature scores in single cells. (F) UMAP representation of single-cell dat at individual days colored by the clusters identified by our balanced clustering routine. (G-H) UMAP representations colored by the gene expression of pluripotent stem cells markers (G) and RPE markers (H) in undifferentiated hESCs and D60 hESC-RPE. (I) Schematic illustrating how the AUC variance evaluation metric behaves in different scenarios (see Figure 1).

**Figure S2.**
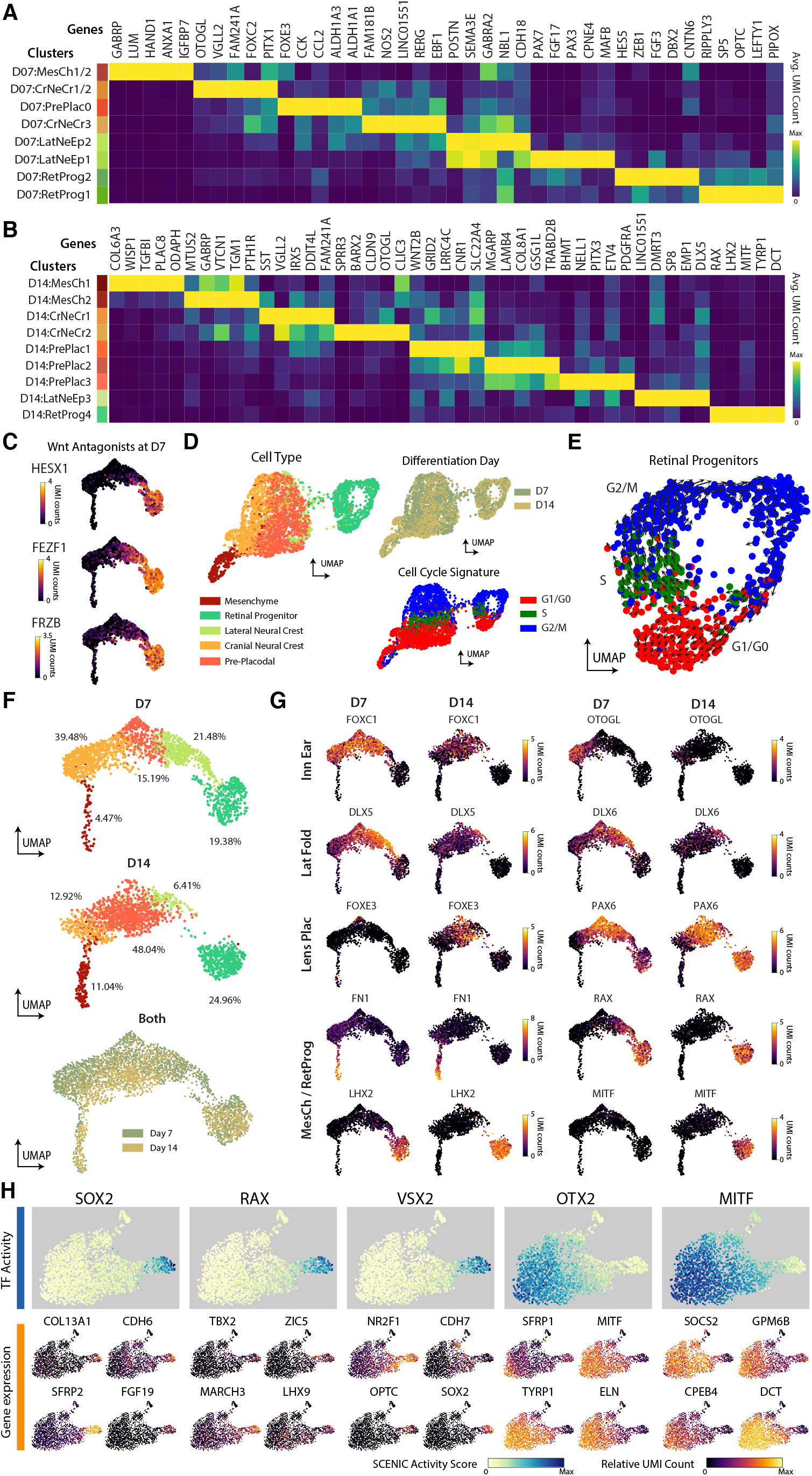
Gene expression characterization and canonical correlation analysis of early hESC-RPE differentiation, Related to Figure 2. (A-B) Heatmap showing the top enriched genes of each cell type cluster at D7 (A) and D14 (B). (C) UMAP embedding (same as (F)) overlayed with the expression of Wnt antagonists *HESX1, FEZF1*, and *FRZB* at D7. (D) UMAP representation of D7 and D14 cell populations analyzed without the removal of cell cycle genes. Cells are colored by cell group, differentiation day, and inferred cell cycle phase. (E) RNA velocity analysis of retinal progenitors at D7 and D14. (F) D7 and D14 cell populations projected on a shared low dimensional subspace using canonical correlation analysis in Seurat (see Methods). (G) UMAP embedding overlayed with the gene expression of key cell type markers for Inner Ear (InnEar), Lateral Fold (LatFold), Lens Placode (LensPlac) and Mesenchymal cells (MesCh)/Retinal Progenitor (RetProg) from D7 to D14. Expression of neural crest inner ear (*FOXC1* and *OTOGL*), and lateral fold (*DLX5* and *DLX6*) decreases from D7 to D14. (H) Transcription factor (TF) activity scores for *SOX2, RAX, VSX2, OTX2*, and *MITF* obtained by SCENIC analysis (top) and the gene expression of the top 4 inferred targets genes of each TF at D30 of differentiation.

**Figure S3.**
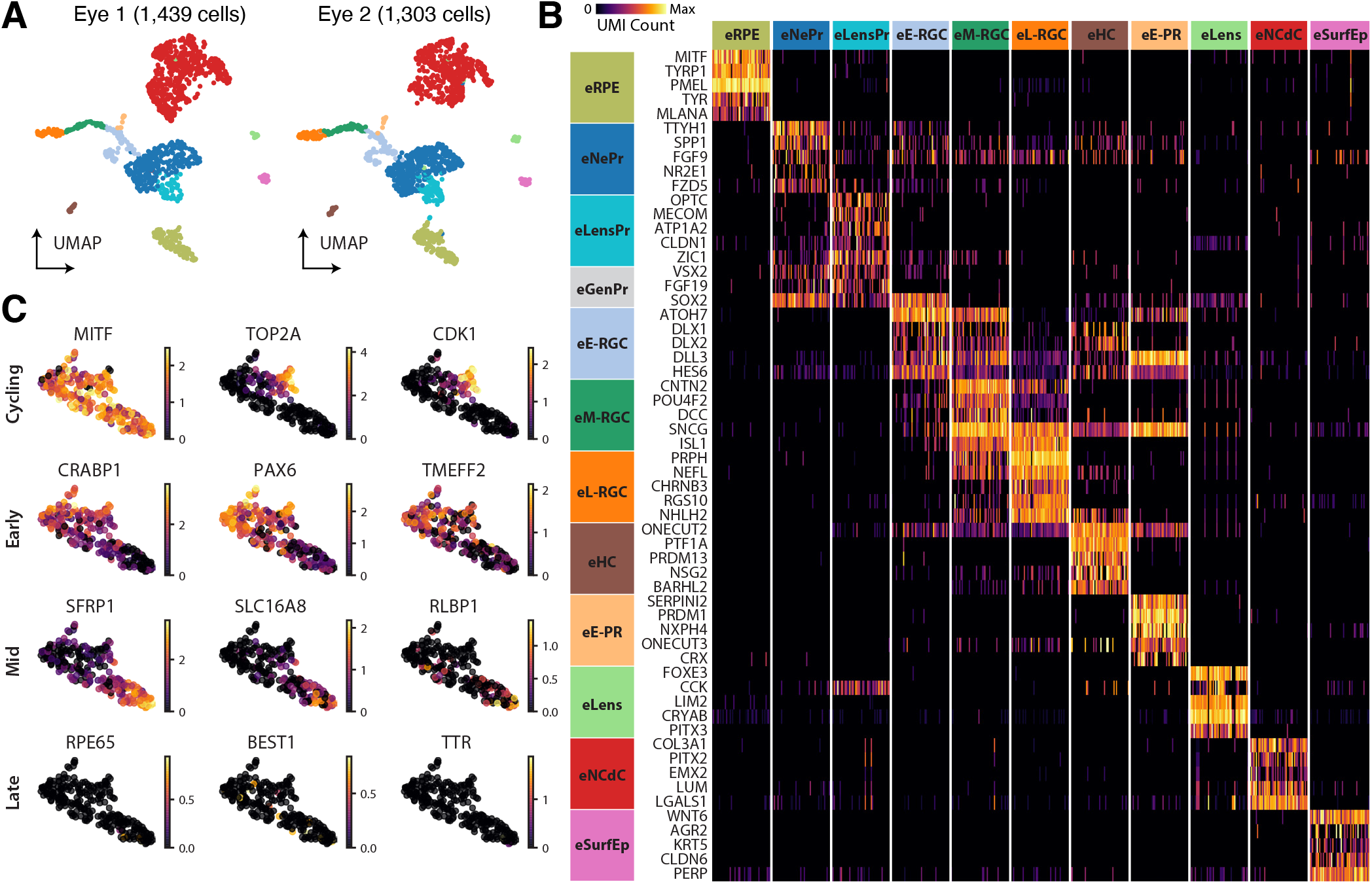
Characterization of the human embryonic eye at week 7.5, Related to Figure 3. (A) UMAP representation of scRNA-seq data from two human embryonic eyes at week 7.5, colored by cell type. (B) Heatmap of normalized enriched gene expression of all cell types at week 7.5. Genes were selected using an enrichment score by cell type in (A): retinal pigment epithelium (eRPE), neural progenitors (eNePr), lens progenitors (eLensPr), early retinal ganglion cells (eE-RGC), mid retinal ganglion cells (eM-RGC), late retinal ganglion cells (eL-RGC), horizontal cells (eHC), early photoreceptors (eE-PR), lens (eLens), neural crest derived cells (eNCdC), and surface epithelium (eSurfEp). eGenPr refers to gene markers shared between the eNePr and eLensPr clusters. (C) Retinal progenitor and RPE log2 normalized gene expression of cycling, early, mid, and late RPE markers in the RPE cell cluster from (A).

**Figure S4.**
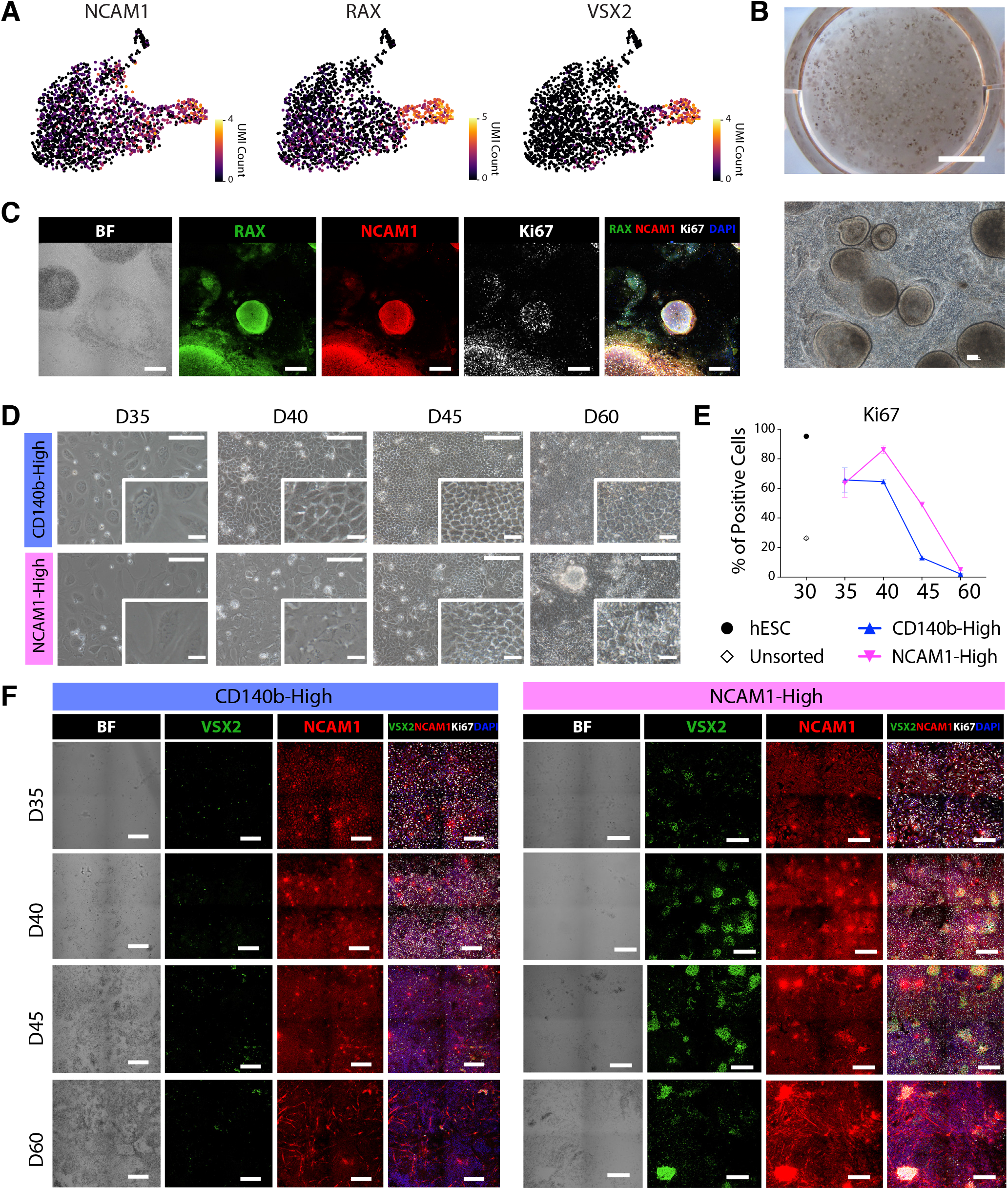
Characterization of the NCAM1 cell sorted cell population, Related to Figure 4. (A) Gene expression of progenitor markers in scRNA-seq hESC-RPE at D30. (B) Camera pictures of hESC-RPE D30 cultures. Scale bars: top 1mm; bottom 100μm. (C) Brightfield and immunofluorescence stainings of hESC-RPE D30 cells showing co-expression of RAX, NCAM1 and Ki67 markers. Scale bars: 200μm. (D) Brightfield images of unsorted, CD140b-High and NCAM1-High populations at D33, D40, D45, and D60 of the differentiation protocol. FACS sorting of the two populations was performed at D30. Scale bars: 100μm; inset 20μm. (E) Graph showing the percentage of positive cells expressing the Ki67 proliferation marker in hESC, unsorted, CD140b-High and NCAM1-High populations at the moment of FACS sorting (D30) and D35, D40, D45, and D60. Bars represent mean +/-SEM from three independent experiments. (F) Brightfield and immunofluorescence images showing expression of VSX2 and NCAM1 in unsorted, CD140b-High and NCAM1-High populations after FACS sorting at differentiation D35, D40, D45, and D60. Scale bars: 200μm.

**Figure S5.**
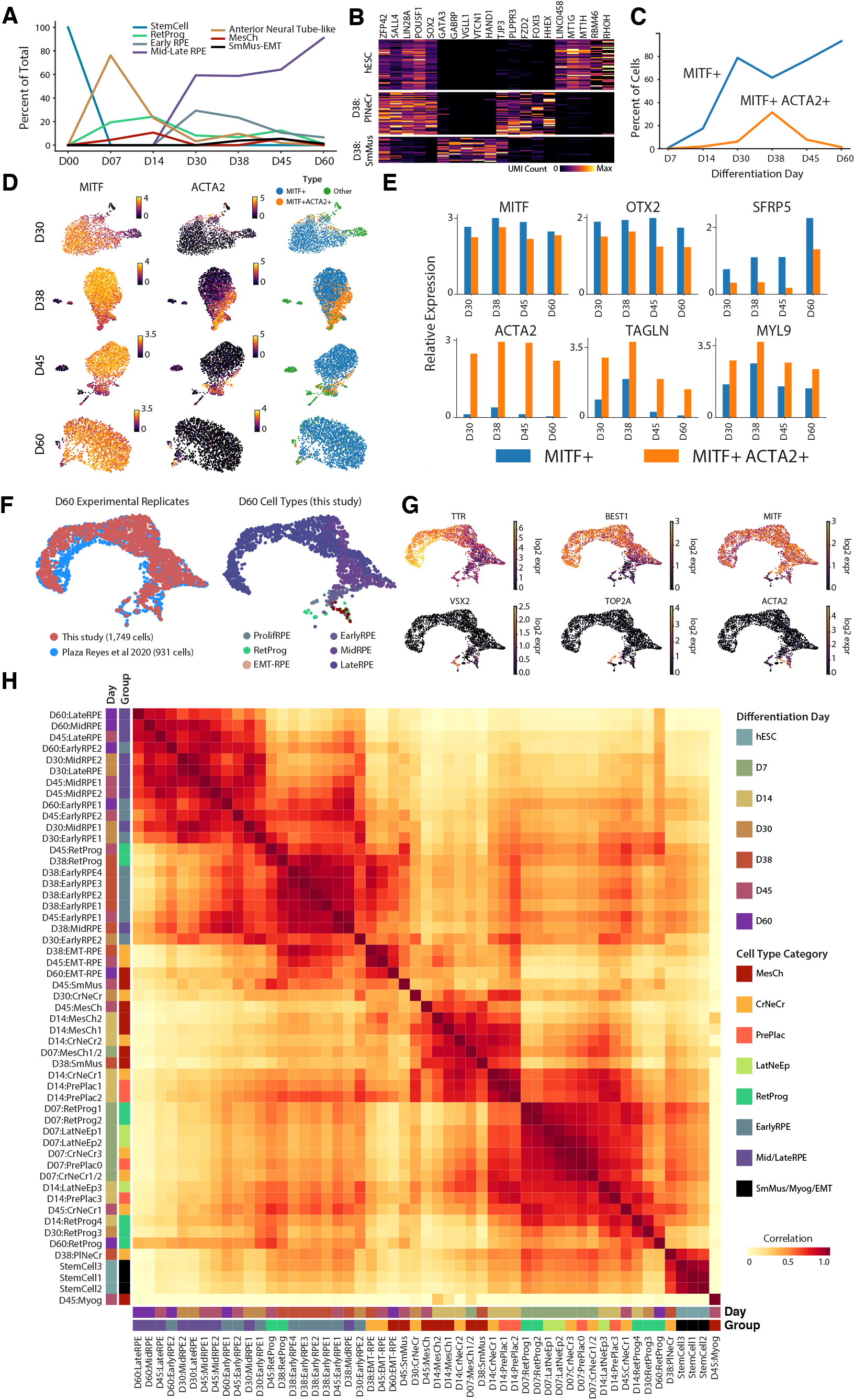
Characterization of late hESE-RPE differentiation and overall gene expression correlation, Related to Figure 6. (A) Line-plot shwoing the percentage of cells assigned to each primary cell type category throughout the entire differentiation time course. (B) Gene expression heatmap showing a comparison of neural crest (NC), and smooth muscle (SM) populations at D38 to the undifferentiated stem cell control. (C) Line-graph showing the change in the proportion of EMT-RPE (MITF+ACTA2+ RPE) and maturing RPE (MITF+) from hESC-RPE D30 to D60. (D) UMAP representation of *MITF* and *ACTA2* gene expression at D30, D38, D45, and D60. Cells are labeled as *MITF+* (maturing RPE), *MITF+ACTA2+* (EMT-RPE), or other (non-RPE cell types). (E) Bar graph showing the gene expression differences between EMT-RPE and maturing RPE. EMT-RPE express EMT markers *ACTA2, TAGLN*, and *MYL9* more highly, whereas RPE markers *MITF, OTX2*, and *SFRP5* are more highly expressed in non-transitioning RPE. (F) CCA integration of 1,749 cells at hESC-RPE D60 from this study with 931 cells at hESC-RPE D60 obtained during a previous experimental replicate from our prior study (Plaza Reyes et al., 2020a). (G) Gene expression representation of D60 replicates for mature RPE (*TTR* and *BEST1*), differentiating RPE (*MITF*), retinal progenitor (*VSX2*), cell cycle (*TOP2A*), and EMT-RPE (*ACTA2*) marker genes. (H) Pearson’s correlation matrix computed using the top 2,000 variable genes among all *in vitro* cell clusters and labeled according to differentiation day (Day) and cluster group (Group).

**Figure S6.**
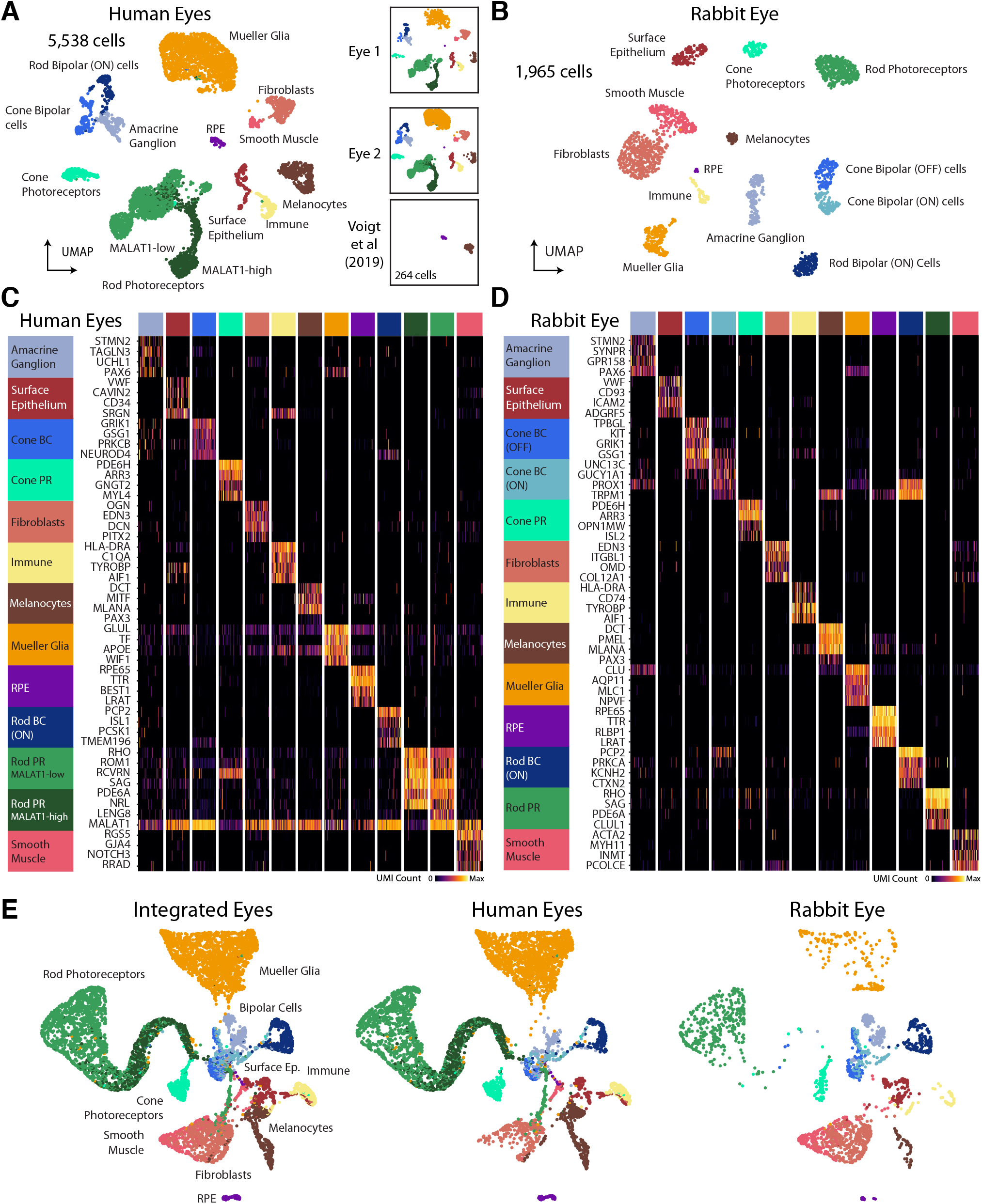
Transcriptional analysis of albino rabbit and human retinas, Related to Figure 7. (A) Annotated UMAP representation of 5,538 human cells from two human eyes (1,564 cells and 3,706 cells), categorized into 13 different cell types. Additional RPEs and melanocytes (264 cells) were re-analyzed and incorporated from Voigt et al, 2019. (B) Annotated UMAP representation of 1,965 rabbit cells categorized into 13 different retina cell types. (C) Heatmap of enriched marker genes for adult eye cell types. BC: bipolar cell. PC: photoreceptor cell. (D) Heatmap of enriched marker genes for rabbit eye cell types. Genes were selected from among the top 20 enriched genes per cluster for (C) and (D). (E) UMAP embedding of the human and rabbit eyes single-cell datasets after CCA integration.

**Figure S7.**
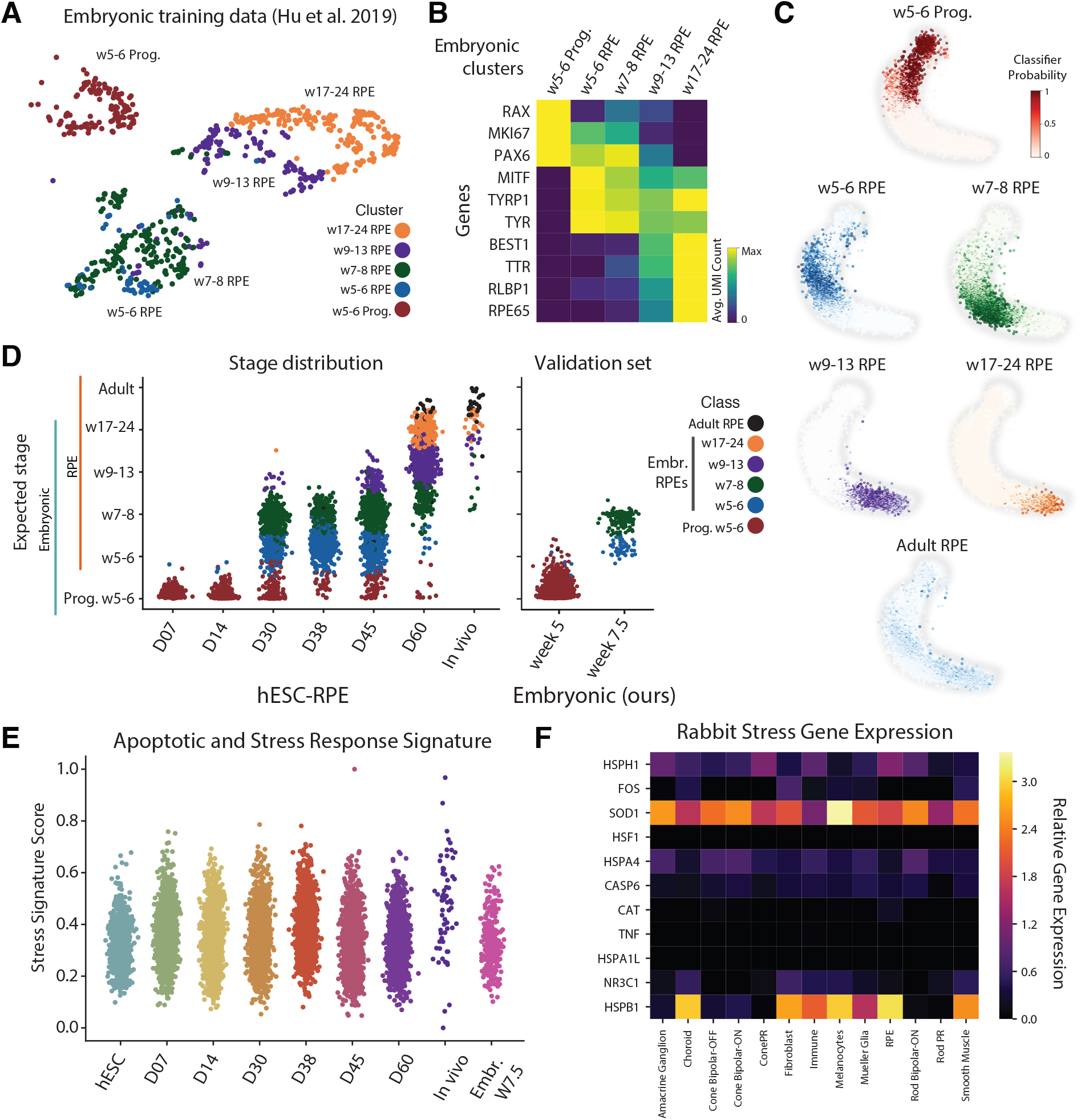
Ordinal classification of *in vitro* hESC-RPE and *in vivo* transplanted RPE with embryonic and adult retinal tissues, Related to Figure 7. (A) UMAP representation of single-cell transcriptomes from Hu et al. 2019 (783 human embryonic cells from various time points) that were used as training dataset for the ordinal classifier (see Figure 7). Colors indicate the five categories defined for the classification task. (B) Heatmap displayng the average gene expression of the uniquely enriched retinal progenitor and RPE marker genes along the five categories. (C) UMAP embedding of the hESC-RPE differentiation data colored by the classification probability y. Intensity of color indicates a higher probability of cell assignment to the respective developmental day. (D) Probability distribution of 7,717 retinal cell classifications from the six differentiation days and *in vivo* transplantation (*In vivo*). Cells were assigned a continuous classification timpoint by computing probability-weighted average of the time points; cells are colored by the category with highest probability. (E) Beaswarm plots showing the signature score for “apoptosis and stress response” in the *in vitro* hESC-RPE cells, *in vivo* grafted, and embryonic W7.5 reference tissues computed using 37 reference genes (see Methods). (F) Heatmap showing the average gene expression of selected stress response genes in rabbit retina post-hESC-RPE subretinal injection.

## SUPPLEMENTAL TABLES

**Table S1.** Summary of the single-cell RNA-sequencing datasets obtained for the characterization of hESC-RPE differentiation protocol and the transcriptomic landscape of the retina, Related to Star Methods. Additional datasets are obtained from the literature and are used and cited appropriately.

**Table S2.** Gene enrichment scores during hESC-RPE pigmentation induction (D7, D14, and D30) by cell type category, Related to Figure 2.

**Table S3.** Gene enrichment scores for embryonic eyes at weeks 5 and 7.5 by cell type category, Related to Figure 3.

**Table S4.** Gene correlation scores with retinal progenitor and neural tube signature during the identification of NCAM1 cell surface marker, Related to Figure 4.

**Table S5.** Gene enrichment scores for adult human and rabbit eyes by cell type category, Related to Figure 7.

## EXTENDED METHODS

### RESOURCE AVAILABILITY

#### Lead Contact

Further information and requests for resources and reagents should be directed to and will be fulfilled by the Lead Contact, Fredrik Lanner.

### MATERIALS AVAILABILITY

This study did not generate new unique reagents.

### DATA AND CODE AVAILABILITY

All scRNA-seq datasets generated for this study are available as in the raw FASTQ format, as generated by CellRanger, at GSE164092. Processed loom files containing spliced and unspliced counts, cell type annotations, UMAP embedding coordinates, and other metadata are also provided. Additionally, we provide the custom reference genome used for human and rabbit pooled analysis. Various lists enriched genes, correlation scores, and other reference data are provided as Supplementary Tables. The external scRNA-seq reference data used in this study can be found at GSE135922 and GSE107618 (Hu et al., 2019; Voigt et al., 2019). Companion source code is available at https://github.com/lamanno-epfl/rpe_differentiation_profiling_code.

### hESC Cell Culture

hESC line HS980 was previously derived and cultured under xeno-free and defined conditions (Rodin et al., 2014) (Swedish Ethical Review Authority: 2011/745:31/3). Donors gave their informed consent for the derivation and subsequent use of hESC lines. The WA09/H9 hESC line was obtained from Wicell and adapted to feeder-free culture on human recombinant LN-521 (hrLN-521, 10 μg/mL, Biolamina). Cells were maintained by clonal propagation on hrLN-521-coated plates in NutriStem hPSC XF medium (Biological Industries) and a 5% CO_2_/5% O_2_ incubator.

Cells were passaged enzymatically at a 1:10 ratio every 5-6 days. Confluent cultures were washed twice with PBS without Ca^2+^ and Mg^2+^ and incubated for 5 min at 37°C, 5% CO_2_/5% O_2_ with TrypLE Select (ThermoFisher Scientific, 12563011). The enzyme was then carefully removed and cells were collected in fresh pre-warmed NutriStem hPSC XF medium by gentle pipetting to obtain a single cell suspension. Cells were centrifuged at 300 g for 4 min, the pellet resuspended in fresh prewarmed NutriStem hPSC XF medium, and plated on a freshly hrLN-521 coated dish. Two days after passage, the medium was replaced with fresh prewarmed NutriStem hPSC XF medium and changed daily.

### hESC-RPE *in vitro* differentiation

A step-by-step protocol describing the differentiation procedure has been reported previously (Plaza Reyes et al., 2020a, 2020b). hESCs were plated at a cell density of 2.4×10^4^ cells/ cm^2^ on hrLN-521 (20 μg/mL) using NutriStem hPSC XF medium. A Rho-kinase inhibitor (Millipore, Y-27632) at a concentration of 10 μM was added during the first 24h, while cells were kept at 37°C, 5% CO_2_/5% O_2_. After 24h, hPSC medium was replaced with differentiation medium NutriStem hPSC XF without bFGF and TGFβ and cells were placed at 37°C, 5% CO_2_/21%O_2_. From day 6 after plating, 100 ng/mL of Activin A (R&D Systems, 338-AC-050) was added to the media. Cells were fed three times a week and kept for 30 days. Monolayers were then trypsinized using TrypLE Select (ThermoFisher Scientific, 12563011) for 10 min at 37°C, 5% CO_2_. The enzyme was carefully removed and the cells were collected in fresh pre-warmed NutriStem hPSC XF medium without bFGF and TGFβ by gentle pipetting to obtain a single cell suspension. Cells were centrifuged at 300 g for 4 min, the pellet was resuspended, passed through a cell strainer (ø 40 μm, VWR, 732-2757) and seeded on laminin coated dishes (hrLN-521 at 20μg/mL) at different cell densities ranging from 1.4×10^6^ to 1.4×10^4^ cells/cm^2^. Replated cells were fed three times a week during the subsequent 30 days with NutriStem hPSC XF medium without bFGF and TGFβ. For hPSC-RPE *in vitro* differentiation in 3D suspension embryoid bodies see our previous publication (Plaza Reyes et al., 2016). Brightfield images were acquired with a Nikon Eclipse TE2000-S microscope and a Canon SX170 IS camera was used to capture pigmentation from the top of the wells.

### Flow Cytometry and Cell Sorting

hPSC-RPE were dissociated into single cells using TrypLE Select (ThermoFisher Scientific, 12563011). Samples were stained with BV421 Mouse Anti-Human CD140b (BD Biosciences 564124, clone [28D4], 10 μg/mL), BB515 Mouse Anti-Human CD56 (BD Biosciences 564489, clone [B159], 2.5 μg/mL), PE Mouse anti-human Ki-67 (Biolegend 350504, clone [Ki-67], 50 μg/mL) conjugated antibodies, diluted in 2% FBS and 1 mM EDTA (ThermoFisher Scientific, 10082147). Cells were incubated with the conjugated antibodies on ice for 30 min. Fluorescence minus one (FMO) controls were included for each condition to identify and gate negative and positive cells. Stained cells were analyzed using a CytoFLEX flow cytometer equipped with 488 nm, 561 nm, 405 nm and 640 nm lasers (Beckman Coulter). Analysis of the data was carried out using FlowJo v.10 software (Tree Star).

Cell sorting was performed on hPSC-RPE cultures after 30 days of differentiation. Cells were incubated with the mentioned conjugated antibodies on ice for 30 min. Fluorescence minus one (FMO) controls were included to identify and gate negative and positive cells. Stained cells were then sorted using a BD FACS Aria Fusion Cell Sorter (BD Bioscience) using FACSDiva Software v8.0.1.

### Neuroretinal progenitor *in vitro* differentiation

Directly after sorting, 68,420 cells/cm2 were plated on matrigel coated plates. Cells were cultured until confluency in P1 media with DMEM/F12 (ThermoFisher Scientific, 11320033) as basal media containing B27 (ThermoFisher Scientific, 17504044), N2 (ThermoFisher Scientific, 17502048), 10ng/mL hDKK1 (R&D Systems, 5439-DK-010), 10ng/ mL mouse Noggin (R&D Systems, 1967-NG-025), 10ng/ mL hIGF-1 (R&D Systems, 291-G1-200), 5ng/mL bFGF (ThermoFisher Scientific, 13256029) and 50 U/mL of Penicillin-Streptomycin (ThermoFisher Scientific, 15140122). Thereafter and until day 40, cells were continued cultured in P1 with the addition of 40ng/mL 3,3’,5-Triiodo-L-thyronine T3 (Sigma, T-074-1ML) and 100uM Taurine (Sigma, T8691-25G). This protocol was based on previous work (Shao et al., 2017).

### qPCR

Total RNA was isolated using the RNeasy Plus Mini Kit and treated with RNase-free DNase (both from Qiagen, 74106 and 79254, respectively). cDNA was synthesized using 1 μg of total RNA in a 20 μL reaction mixture, containing random hexamers and Superscript III reverse transcriptase (ThermoFisher Scientific, 18080085), according to the manufacturer’s instructions. Taq-polymerase together with Taqman probes (ThermoFisher Scientific) for *MITF* (Hs01117294_ m1), *BEST1* (Hs00188249_m1), *RPE65* (Hs01071462_ m1), *TYR* (Hs00165976_m1), *SIX6* (HS00201310_m1), *PAX6* (HS01088114_m1), *VSX2* (HS01584046_m1), *RAX* (HS00429459_m1), *SOX2* (HS04234836_s1), and *GAPDH* (4333764F) were used. Samples were subjected to the real-time PCR amplification protocol on StepOneTM real-time PCR System (Applied Biosystems). Three independent experiments were performed for every condition and technical duplicates were carried for each reaction. Results are presented as mean ± SEM (standard error of the mean).

### Immunocytofluorescence

Protein expression was assessed with immunofluorescence. Cells were fixed with 4% methanol-free formaldehyde (VWR, FFCHFF22023000) at room temperature for 10 min, followed by permeabilization with 0.3% Triton X-100 (Sigma, T9284) in Dulbeccos phosphate-buffered saline (D-PBS, ThermoFisher Scientific, 14190094) for 10 min and blocking with 4% fetal bovine serum (FBS, ThermoFisher Scientific, 10082147) and 0.1% Tween-20 (Sigma, P9416) in DPBS for 1 hour. Primary antibodies were diluted to the specified concentrations in 4% FBS, 0.1% Tween-20, DPBS solution: VSX2/Chx10 (1:50, Santa Cruz Biotechnology sc-365519, clone [E-12]), Ki67 (1:400, Cell Signaling Technology 9027, clone [D2H10]), Bestrophin 1 (BEST-1) (1:100, Millipore MAB5466), RAX (10ug/mL, Novusbio H00030062-M02, clone [4F4]), PDGFRB (CD140b) (1:100, BD Biosciences, 558820, clone [28D4]) and NCAM1 (CD56) (1:100, BD Biosciences, 555513, clone [B159]). The primary antibodies were incubated overnight at 4°C followed by 2 hours incubation at room temperature with secondary antibodies: donkey anti-mouse IgG (H+L) Alexa Fluor 488, donkey anti-mouse IgG Alexa Fluor (H+L) 555, donkey anti-mouse IgG Alexa Fluor (H+L) 647, donkey anti-rabbit IgG (H+L) Alexa Fluor 647 (all of them from ThermoFisher Scientific, A21202, A31570, A31571, A31573, respectively) diluted 1:1,000 in 4% FBS, 0.1% Tween-20, D-PBS solution. Nuclei were stained with Hoechst 33342 (1:1,000, Invitrogen H3570). Images were acquired with Zeiss LSM710-NLO point scanning confocal microscope. Post-acquisition analysis of the pictures was performed using ImageJ v2.0 software.

### Histology and Tissue Immunostaining

Immediately after euthanasia by intravenous injection of 100 mg/kg pentobarbital (Allfatal vet. 100 mg/mL, Omnidea), the eyes were enucleated and the bleb injection area marked with green Tissue Marking Dye (TMD; Histolab Products AB, 02199). An intravitreal injection of 100 μL fixing solution (FS) consisting of 4% buffered formaldehyde (Histolab Products AB, 02175) was performed before fixation in FS for 24-48 hours and embedding in paraffin. Four-μm serial sections were produced through the TMD-labeled area. For immunostaining, slides were deparaffinized in xylene, dehydrated in graded alcohols, and rinsed with ddH2O and Tris Buffered Saline (TBS, Sigma, 93352, pH 7.6). Antigen retrieval was achieved in 10 mM citrate buffer (trisodium citrate dihydrate, Sigma, S1804, pH 6.0) with 1:2000 Tween-20 (Sigma, P9416) at 96°C for 30 min, followed by 30 min cooling at room temperature. Slides were washed with TBS and blocked for 30 min with 10% Normal Donkey Serum (Abcam, ab138579) diluted in TBS containing 5% (w/v) IgG and protease-free bovine serum albumin (Jackson Immunoresearch, 001-000-162) in a humidified chamber. Primary antibodies diluted in the blocking buffer were incubated overnight at 4°C: human nuclear mitotic apparatus protein (NuMA) (1:200, Abcam ab84680), BEST-1 (1:200, Millipore MAB5466). Secondary antibodies (donkey anti-rabbit IgG (H+L) Alexa Fluor 555 A31572 and donkey anti-mouse IgG (H+L) Alexa Fluor 647 A31571, both from ThermoFisher Scientific) diluted 1:200 in blocking buffer, were incubated 1 hour at room temperature. Sections were mounted with vector vectashield with DAPI mounting medium (Vector Laboratories, H-1200-10) under a 24×50 mm coverslip. Images were taken with an Olympus IX81 fluorescence inverted microscope or Zeiss LSM710-NLO point scanning confocal microscope. Post-acquisition analysis was performed using ImageJ v2.0 software.

### Subretinal Injections

hESC-RPE monolayers were washed with DPBS (ThermoFisher Scientific, 14190-094), incubated with TrypLE (ThermoFisher Scientific, 12563-011) and dissociated to single cell suspension as described above. Cells were counted in a Neubauer hemocytometer (VWR, 631-0925) chamber using 0.4% trypan blue (ThermoFisher Scientific, 15250061), centrifuged at 300g for 4 min, and the cell pellet was resuspended in freshly filter-sterilized DPBS (ThermoFisher Scientific, 14190-094) to a final concentration of 1000 cells/μL. The cell suspension was then aseptically aliquoted into 600 μL units and kept on ice until surgery.

After approval by the Northern Stockholm Animal Experimental Ethics Committee (DNR N56/15), two female New Zealand white albino rabbits (provided by the Lidköpings rabbit farm, Lidköping, Sweden) aged 5 months and weighing 3.5 to 4.0 kg were used in this study. All experiments were conducted in accordance with the Statement for the Use of Animals in Ophthalmic and Vision Research.

Animals were put under general anesthesia by intramuscular administration of 35 mg/kg bw (body weight) ketamine (Ketaminol 100 mg/mL, Intervet, 511519) and 5 mg/kg xylazine (Rompun vet. 20 mg/mL, Bayer Animal Health, 22545), and the pupils were dilated with a mix of 0.75% cyclopentolate / 2.5% phenylephrine (APL, 321968). Microsurgeries were performed on both eyes using a 2-port 25G transvitreal pars plana technique (Alcon Nordic A/S, 8065751448). 25G trocars were inserted 1 mm from the limbus and an infusion cannula was connected to the lower temporal trocar. The cell suspension was drawn into a 1 mL syringe connected to an extension tube and a 38G polytip cannula (MedOne Surgical Inc, 3219 and 3223). Without infusion or prior vitrectomy the cannula was inserted through the upper temporal trocar. After proper tip positioning, ascertained by a focal whitening of the retina, 50 μL of cell suspension (equivalent to 50,000 cells) was injected slowly subretinally, approximately 6 mm below the inferior margin of the optic nerve head, forming a uniform bleb that was clearly visible under the operating microscope. To minimize reflux, the tip was maintained within the bleb during the injection. After instrument removal light pressure was applied to the self-sealing suture-less sclerotomies. 2 mg (100 μL) of intravitreal triamcinolone (Triescence 40 mg/mL, Alcon Nordic A/S, 412915) was administered a day prior to the surgery, and no post-surgical antibiotics were given.

### Retinal tissue dissociation of human adult, fetal and rabbit eyes with hESC-RPE for single-cell RNA sequencing

Human post-mortem research-consented donor eyes were obtained from the cornea bank at St. Erik Eye Hospital, Stockholm, Sweden. The use of human tissue was in accordance with the tenets of the Declaration of Helsinki and was approved by the Swedish Ethical Review Authority (EPN, #2019-02032) for the use of human donor material for research. Donors did not present any clinical diagnosis of ocular disease, and samples were anonymized and processed under the general data protection regulation. Two human eyes from the same donor were used (45-year-old male, 32 hours post-mortem). The lens was dissected out and the rest of the retina, except the sclera, was chopped in several small pieces mixed together in 500 uL of digestion buffer (described below). Two embryonic eyes from the same embryo were used from a 7.5 post-conception week embryo. The optic cups were dissected out and chopped in several small pieces to facilitate dissociation in 500 uL of digestion buffer (described below). Donors (deceased, family for adult human eyes, or couples for the embryos) gave their informed consent for the donation and subsequent use for research purposes. The embryonic eyes were acquired from a clinical routine abortion after informed consent by the pregnant woman, in accordance with permissions from the regional ethical review board and the Swedish National Board of Health and Welfare (“Socialstyrelsen” #8.1-11692/2019) and the Swedish Ethical Review Authority (“EPN” #2007/1477-31/3. Two rabbit eyes (from different animals) with 30-day integrated hESC-RPE were enucleated and pigmented areas including neuroretina, choroid and RPE layer was dissected out, trimmed and mixed together in 500 uL of digestion buffer. Digestion buffer consisted of: 2mg/ml collagenase IV (ThermoFisher Scientific, 17104019), 120 U/μl DNase I (NEB, Sigma, 4536282001), and 1mg/ml papain (Sigma, 10108014001) in PBS. Eppendorfs containing the samples were rotated and incubated at 37°C on a thermocycler at 300g for 25 min until samples were homogenized. Samples were pipetted every 5 min to digest the tissue sample into single cells. Digestion was stopped by adding equivalent volume of 10% fetal bovine serum (ThermoFisher Scientific, 10082147) in PBS, the samples were filtered using a 30μm MACS Smart Strainer (Miltenyi, 130-098-458) followed by Dead Cell Removal kit (Miltenyi, 130-090-101) to remove dead cells and debris. At this stage, one of the rabbit eye cell samples was stained with mouse anti-human HLA-ABC-FITC (1:20, BD Biosciences, 555552, clone [G46-2.6]), and anti-human HLA-ABC-positive cells were FACS-sorted as specified above, collected and resuspended to 1000 cells/ uL in 1% BSA (Sigma, A1470-10g) in PBS for further scR-NA-sequencing. Otherwise, cells were finally resuspended to 1000 cells/uL in 1% BSA in PBS prior to scRNA-sequencing. Acquisition of all primary human tissue samples from two embryonic eyes at week 5 (Carnegie Stage 13) was approved by the UCSF Human Gamete, Embryo and Stem Cell Research Committee (GESCR, approval 10-03379 and 10-05113). All experiments were performed in accordance with protocol guidelines. Informed consent was obtained before sample collection and use for this study. First-trimester human samples were collected from elective pregnancy terminations through the Human Developmental Biology Resource (HDBR), staged using crown-rump length (CRL) and shipped overnight on ice in Rosewell Park Memorial Institute (RPMI) media. Dissections were based upon anatomical landmarks, and dissociations were performed using papain (Worthington). Samples were incubated in papain for 20-30 minutes and triturated manually into a single cell suspension. The samples were filtered for remaining debri and moved to PBS with 0.05% BSA to be captured by 10X Genomics Chromium RNA capture version 2. Library preparation was performed based upon manufacturer’s instructions, and sequencing was performed on a NovaSeq S4 lane.

### Single-cell RNA sequencing

Cells were transported at 4°C to the Eukaryotic Single Cell Genomics Facility (ESCG, SciLifeLab, Stockholm, Sweden) where a 3’ cDNA library was prepared for single cell RNA sequencing (scRNA-seq) using the 10X Genomics platform and NovaSeq 6000 software. Cell Ranger 2.1.1 was used to convert Illumina base call files to FASTQ format. Cell Ranger 3.1.0 was used to map FASTQ sequencing reads to the human GRCh38 reference transcriptome with the STAR aligner and to generate feature-barcode count matrices. The velocyto run10x command was used to produce loom files containing spliced and unspliced RNA count matrices and most analyses (described below) were performed using ve-locyto and scanpy in Jupyter Notebooks (La Manno et al., 2018; Wolf et al., 2018).

### scRNA-seq processing for hESC-RPE *in vitro* differentiation

The initial number of barcoded cells was 14,997 cells (1,790 hESCs, 2,280 cells at D7, 2,451 cells at D14, 2,387 cells at D30, 2,075 cells at D38, 1,939 cells at D45, and 2,075 cells at D60). As a quality-control, only cells with spliced UMIs (>8,000 and <75,000), unspliced UMIs (>2,500 and <30,000), uniquely expressed genes (>2,500) and a low percentage of mitochondrial reads (<15%) were retained, resulting in 11,791 single cells (1,016 hESCs, 1,811 cells at D7, 1,872 cells at D14, 1,852 cells at D30, 1,784 cells at D38, 1,707 cells at D45, and 1,749 cells at D60) (Table S1). Cell cycle phase assignments were inferred using scanpy.score_genes_cell_ cycle on unnormalized spliced counts and with published signature genes (Satija et al., 2015; Wolf et al., 2018). The cell cycle signature was then disentangled from downstream analysis by using a Wilcoxon rank sum test to find and exclude the top 1,500 enriched genes by phase (G1/G0, S, or G2/M) from feature selection. 2,515 genes cycle-enriched in >=4/7 time points were excluded. Feature selection and size normalization were performed using velocyto.cv_vs_mean and velocyto.normalize. PCA, UMAP, and Louvain clustering were performed with standard parameters in scanpy and velocyto using spliced counts (2,000 selected genes, 20 PCs, 50 nearest neighbors). Global dimensionality reduction of the entire dataset was performed using a profile of pluripotency (6 genes: *LIN28A, NANOG, POU5F1, SALL4, SOX2, ZFP42*), retinal progenitor (20 genes: *BMP4, FGF3, FGF8, LHX2, NCAM1, OPTC, OTX1, OTX2, PAX2, PAX6, RAX, SIX3, SIX6, TBX2, TBX3, TBX5, VAX2, VSX2, ZIC1, ZIC2*) and RPE (16 genes: *BEST1, ELN, IGFBP5, MITF, PMEL, RGR, RLBP1, RPE65, SERPINF1, SLC7A8, TMEFF2, TRPM1, TRPM3, TTR, TYR, TYRP1*) marker genes, and the first two principal components derived from normalized spliced counts were visualized (Bosze et al., 2020; Fuhrmann, 2010; Schmitt et al., 2009). Signature score for apoptosis and stress response were computed using the following 37 genes: *CASP8, CASP10, E2F5, CASP2, PTEN, CASP7, CDKN2A, E2F3, E2F4, CASP6, TP53, E2F1, CDKN1B, CDKN1A, E2F2, CASP3, FHIT, CASP9, HSPH1 GPX3, SOCS3, FOS, HSPA1A, SOD2, SOD1, HSF1, SOD3, HSPA1B, HSPA4, CASP6, GPX4, CAT, TNF, IL6, HSPA1L, NR3C1, HSPB1*. Signature scores for three cell types identities were computed across all cells by scaling each gene (above) between 0 and 1 and summing total scaled expression for all included markers.

To ensure equalized cluster resolution across individual differentiation days, each time point was initially over-clustered using the resolution parameter of Louvain clustering. Using the sklearn Logistic Regression classifier (3-fold cross validation), low-scoring clusters were recursively merged until the classifier achieved greater than 80% prediction accuracy, on the test set, for each cluster and a change in minimum accuracy of less than 5% between the iterations. The result was 52 clusters with >5 cells (56 in total) across seven differentiation days. The percent variance contained within the clusters was calculated as follows: first, we found the Euclidean distance between cells and cluster centroids with sklearn.NearestCentroid and scipy.spatial.distance. The total pairwise-cluster sum of squares and total within-cluster sum of squares were then computed; lastly the difference (between-cluster sum of squares) was divided by the pairwise-cluster sum of squares to obtain the percent variance. Subsequently, clusters were independently assessed and merged if necessary for downstream gene enrichment analysis, leading to annotation of 11 primary and 53 secondary cell type cluster categories. SCENIC transcription factor regulon analysis was performed using pyscenic and default parameters as described in the original study (Aibar et al., 2017).

### scRNA-seq processing for human reference tissues

From the embryonic optic vesicle at week 5 (Carnegie stage 13), all 2,637 single cells were analyzed using standard parameters in velocyto and scanpy (800 CV-mean genes, 20 PCs, 20 nearest neighbors) (Table S1). Gene enrichment scores were computed and applied to annotate the following cell clusters: Neural Retina (956 cells), Mesenchymal (471 cells), Retinal Pigment Epithelium (452 cells), Neural Tube (424 cells), Optic stalk (179 cells), Smooth Muscle (40 cells), and Immune cells (13 cells). A cluster of blood cell contaminants (102 cells) was excluded from downstream analyses.

3,054 unfiltered single cells from two human embryonic eyes at week 7.5 (1,577 from eye 1 and 1,477 from eye 2) were analyzed using velocyto and scanpy. Cell Ranger 2.1.1 was used to map FASTQ sequencing reads to the human GRCh38 reference transcriptome with the STAR aligner and to generate feature-barcode count matrices. As a quality-control, only cells with UMIs (>3,000 and <50,000), uniquely expressed genes (>3,000 and <7,000) and a low percentage of mitochondrial reads (<10%) were retained, resulting in 2,742 single cells (1,439 from eye 1 and 1,303 from eye 2) (Table S1). Due to minimal batch effects between embryonic eyes, cv_vs_mean selection (3,000 genes), dimensionality reduction, and other downstream analyses were jointly performed without batch effect correction (3,000 CV-mean genes, 25 PCs, 50 nearest neighbors). Cells were assigned to clusters annotated using gene enrichment as follows: Neural Crest Derived (1,078 cells), Neural Progenitors (706 cells), Retinal Pigment Epithelium (258 cells), Early Ganglion (165 cells), Lens Progenitors (150 cells), Mid Ganglion (104 cells), Late Ganglion (88 cells), Surface Epithelium (53 cells), Horizontal Cells (36 cells), Lens (35 cells), and Early Photoreceptors (22 cells). Blood cell contaminant was excluded (47 cells). The neural crest derived cell cluster was re-clustered a second time, resulting in 7 subclusters.

6,215 unfiltered single cells from two human adult eyes (3,708 from eye 1 and 1,566 from eye 2) were analyzed using velocyto and scanpy. As a quality-control, only cells with UMIs (>500 and <30,000), uniquely expressed genes (>100) and a low percentage of mitochondrial reads (<10%) were retained, resulting in 5,538 single cells (4,420 cells from eye 1 and 1,792 from eye 2) (Table S1). Due to the limited batch effect between eyes, downstream analyses were performed on the joined expression matrix. To strengthen our reference and ensure accurate capture of all RPE cells, a subset of RPE and melanocyte cells from scRNA-seq in a prior study were included in downstream analysis (reprocessed and filtered according to identical parameters) (Voigt et al., 2019). These 263 additional cells were obtained from 8 samples by performing a clustering of the published, reprocessed counts and selecting cells expressing melanocyte marker *MLANA* or RPE marker *RPE65* (GEO accession number: GSE135922). Adult retinal cell types were annotated using known marker genes and genes with high enrichment scores as follows: Muller Glia (1,646 cells), Rod Photoreceptors (MALAT1-lo) (1,323 cells), Rod Photoreceptors (MALAT1-hi) (747 cells), Melanocytes (364 cells), Fibroblast (267 cells), Cone Photoreceptors (218 cells), Cone Bipolar (195 cells), Amacrine Ganglion (181 cells), Rod Bipolar-ON (169 cells), Surface Epithelium (161 cells), Immune (136 cells), Retinal Pigment Epithelium (71 cells), and Smooth Muscle (60 cells).

### scRNA-seq processing for NCAM1-High-derived *in vitro* neuronal differentiation

NCAM1-High-derived cells from our alternative neuronal differentiation protocol were analyzed using velocyto and scanpy on 2,638 unfiltered single cells. Cells with UMIs (>6,000 and <100,000) and a low percentage of mitochondrial reads (<10%) were retained during filtering. After dimensionality reduction and UMAP analysis (3,000 CV-mean genes, 20 PCs, 20 nearest neighbors), as well as the removal of a cluster cell contaminant, we obtained 980 single cells (Table S1) of the following clusters, upon which we performed gene enrichment: Mesenchymal (447 cells), Retinal Progenitors (134 cells), Retinal Pigment Epithelium (121 cells), Early Neuroblasts (102 cells), Neuroblasts (77 cells), Glutamatergic Neurons (46 cells), Gabaergic Neurons (25 cells), Surface Epithelium (23 cells), and Lens (5 cells).

An integrated subspace was found between NCAM1-High-derived cells (980 cells) and human embryonic eyes at week 7.5 (2,742 cells, see above) using CCA integration and 2,000 CV-mean genes for each dataset (3,063 unique genes in total). Gene enrichment scores were computed to identify unique and shared markers between the various populations, including overlapping lens and surface epithelium clusters. RNA velocity was performed on NCAM1-High-derived neurons (Early Neruoblast, Neuroblast, and Glutamatergic Neuron clusters) and embryonic retinal ganglion cells (Early, Mid, and Late Ganglion clusters) with velocyto.

To compare the differentiation programs of NCAM1-High-derived *in vitro* neurons and embryonic retinal ganglion neurons, we fit a principal curve to the two-dimensional UMAP embedding and identified a pseudotemporal cell ordering with which we could perform pseudotime gene expression alignment analysis. Principal curve code can be found at: https://github.com/lamanno-epfl/rpe_differentiation_profiling_code and is adapted from the original publication (Hastie and Stuetzle, 1989). Along each pseudotime, a set of five normally distributed and spaced curves were generated using scipy.stats.norm.pdf to mimic different possible expression patterns along the differentiation trajectory (i.e. early downregulation, upregulation midway through the pseudotime followed by downregulation, upregulation at the end of the pseudotime, etc.). Pearson’s correlation coefficients were computed between genes and each normally distributed peak in order to identify genes with a distinct temporal upregulation and/or downregulation. For *in vitro* neurons and embryonic neurons, the top 50 genes for each of five peaks were combined and compared, enabling us to visualize genes with a similar expression behavior along both pseudotimes as well as genes with a distinct expression behavior unique to either *in vitro* neurons or embryonic neurons alone.

### scRNA-seq processing for hESC-RPE *in vivo* rabbit subretinal injection

Unlike the other scRNA-seq data in this study, feature-barcode count matrices for hESCs isolated from rabbit retina were generated using a custom-built hybrid reference transcriptome combining both the human GRCh38 reference and the rabbit reference (Oryctolagus cuniculus 2.0.99, EMBL-EBI). FASTQ files were mapped to this hybrid reference with Cell Ranger 3.1.0 to generate count matrices. Reads were exclusively assigned either to a human or rabbit transcript. The percentage of UMIs per cell assigned to the human or rabbit reference was then computed. Human cells were retained if >80% reads mapped to the human genome. Rabbit cells were retained if >95% reads mapped to the rabbit genome. The resulting number of cells were 24 human cells from the unsorted approach (0.8% of total, Rabbit1, unsorted) and 61 human cells by sorted approach (4.3% of total, Rabbit 2, sorted). As a control, feature-barcode count matrices were also created by aligning FASTQ files solely to the human or rabbit genome with Cell Ranger.

For analysis of human cells, cells with > 30% reads mapped to mitochondrial genes were removed, resulting in a total of 65 single cells (Table S1). Counts were size normalized in scanpy and gene feature selection was performed to select 100 variable genes using cv_vs_mean. Following log-transformation of counts, expression of RPE marker genes and other highly enriched genes was assessed. Differentiation expression analysis was performed by computing fold change on log2 normalized counts.

For analysis of rabbit retina, only cells obtained from the unsorted approach were used in order to avoid any potential cell-type sorting bias. Cells were filtered to retain those with a certain number of UMIs (>2,000 to <50,000), uniquely expressed genes (>600 to <6,000) and mitochondrial reads (<10%). The total number of cells after filtering was 1,965 cells. Dimensionality reduction and clustering analysis was performed using 3,000 cv_vs_mean selected genes and the workflow described above. Cell types were annotated using known marker genes and genes with high enrichment scores as follows: Fibroblast (408 cells), Rod Photoreceptors (338 cells), Smooth Muscles (200 cells), Muller Glia (182 cells), Amacrine Ganglion (156 cells), Rod Bipolar-ON (139 cells), Surface Epithelium (125 cells), Cone Bipolar-OFF (119 cells), Cone Bipolar-ON (105 cells), Cone Photoreceptors (89 cells), Immune (54 cells), Melanocytes (41 cells), and Retinal Pigment Epithelium (9 cells). For integration of the human adult eye and rabbit eye, anchors were found only between genes with shared nomenclature between the two species’ genomes (9,889 genes) and the top enriched 2,000 genes (2,979 genes in total).

### Gene enrichment and signature score analysis

Throughout this study, all marker genes for visualized cell type clusters were selected from among the top 50 ranked genes calculated with unnormalized counts and a gene enrichment score. For each gene, the mean value per cluster was obtained and scaled by the number of cells per cluster with non-zero counts of the given gene. An gene-wise enrichment score was then computed by comparing the means and the fraction of non-zero values among all clusters.

### Cell heterogeneity estimation

When decomposing the covariance of a heterogeneous cell population, the largest axes of variation typically explain cell type variability, whereas the smaller axes describe subtle phenotypic variation and uncorrelated noise accumulating on the remaining principal components. We deduced that the area under the curve (AUC) of cumulative principal component variance would serve as an insightful metric to assess the overall heterogeneity at each differentiation day. For each time point, cv_vs_mean feature selected genes with a score greater than 0.30 were collected, and a union of genes from all time points were retained and size normalized. 800 cells and half the total number of genes were randomly sub-sampled from each time point, without replacement, and PCA analysis was performed. AUC of the cumulative explained variance ratio for all principal components was calculated using numpy.trapz. Cells and genes were randomly subsampled for 1000 iterations to obtain a distribution of AUCs at each differentiation day. As a greater deviation from the variance due to biological noise results in a greater AUC of cumulative explained variance, the AUC can therefore be used as a proxy for assessing the overall dataset heterogeneity. Statistical significance between time points was computed using a bootstrap paired t-test.

### Canonical correlation analysis and data integration

Canonical correlation analysis was performed in all cases using a Python adaptation of the Seurat CCA method and is available at the following GitHub repository: https://github.com/lamanno-epfl/rpe_differentiation_profiling_code (Butler et al., 2018). For integration of the *in vitro* hESC-RPE D7 and D14 time points, both with and without the cell cycle regressed out, anchors were found using the top enriched 1,000 genes at each time point obtained with cv_vs_mean in velocyto. For integration of the human embryonic week 7.5 and NCAM1-High-derived neuronal hESCs, anchors were found using the top enriched 1,000 genes.

### RPE correlation analysis

To identify genes strongly linked to a progenitor fate among *in vitro* hESC-RPE D30 cells, Pearson’s correlation coefficients were calculated between log-normalized gene counts and either an RPE signature score (16 genes, described above) or a neural tube signature (17 genes: *WNT6, COL17A1, CDH1, TP63, KRT19, KRT17, CRABP2, COL3A1, CYP26A1, FOXC1, HAND1, SEMA3D, SOX17, DLX2, DLX3, DLX4, PDGFRA*). For the RPE correlation score, D30 cells from RetProg and RPE secondary clusters were used and for the neural tube correlation score, D30 cells from RPE and CrNeCr secondary clusters were used. Genes with an average spliced expression <0.5 as well as genes included in the signature scores themselves were excluded, resulting in the calculation of RPE signature correlation coefficients for 5,214 genes and of neural signature correlation coefficients for 4,664 genes. Both correlation coefficients were then averaged to obtain the top genes anticorrelated to both signature and therefore most indicative of a cell progenitor status. The computed correlation coefficients are available in Table S3.

### RNA velocity analysis

RNA velocity analysis was performed using velocyto and scvelo, for dimensionality reduction, KNN smoothing, and gamma fitting we used default parameters unless specified (Bergen et al., 2019; La Manno et al., 2018). For embryonic retinal ganglion neurons and NCAM1-High-derived neuronal hESCs, the steady-state implementation of RNA velocity was used (2,000 enriched genes, 20 PCs, 20 nearest neighbors). For estimating RNA velocity of selected *in vitro* hESC-RPE D60 cells, 1,000 CV-mean genes were selected for velocity estimation. To ensure capture of the appropriate genes, we selected genes with a highly coordinated velocity using a velocity coordination function adapted from velocyto and available at: https://github.com/lamanno-epfl/rpe_differentiation_profiling_code.

### Cell type similarity analysis

Correlation analysis was performed as follows: using the 52 sub-cluster assignments and 1,000 cv_vs_mean selected genes (cell cycle enriched genes were previously excluded and 200 cells from each sub-cluster were randomly selected, with replacement, for uniformity), correlation distances between mean expression by cluster were computed using scipy cdist and visualized using seaborn.

### Ordinal classifier

In order to assign *in vitro* RPE cells to an embryonic reference, we constructed an ordinal classifier function in Python and trained it using scRNA-seq data of embryonic RPE from weeks 5 to 24 of development from a prior study (Hu et al., 2019). This classifier is based on prior work and relies on training a series of sequential classifiers representing temporally ordered stages of development (Frank and Hall, 2001). Sequential classes were designed by grouping RPE scRNA-seq gene expression training data by similar embryonic stages (weeks 5-6, week 7-9, weeks 11-13, weeks 17-24, and adult RPE). The implementation is available at https://github.com/lamanno-epfl/rpe_differentiation_pro-filing_code.

### Bioinformatics software

All analysis was performed using Python 3.7.4. The following modules were used: jupyterlab 1.1.4, loompy 3.0.6, louvain 0.6.1, matplotlib 3.3.1, numpy 1.19.1, pandas 0.25.1, pyscenic 0.10.4, python-igraph 0.7.1, python-louvain 0.13, scanpy 1.4.5, scikit-learn 0.23.2, scipy 1.5.2, scvelo 0.1.25, seaborn 0.9.0, umap-learn 0.4.0, velocyto 0.17.17.

### Statistical Analysis

For statistical analysis, one-way ANOVA and posthoc multiple comparisons using Tukey’s test correction wereperformed to assess the *in vitro* differences of the sorted (NCAM1-High, CD140b-High) and unsorted D30 cells in TEER and PEDF secretion assays. Standard error of the mean and standard deviation calculations for all single cell analyses were performed using the numpy package in Python.

## REFERENCES

Adler, R., and Canto-Soler, M.V. (2007). Molecular mechanisms of op

Aibar, S., González-Blas, C.B., Moerman, T., Huynh-Thu, V.A., Imrichova, H., Hulselmans, G., Rambow, F., Marine, J.-C., Geurts, P., Aerts, J., et al. (2017). SCENIC: single-cell regulatory network inference and clustering. Nat. Methods 14, 1083–1086.

Ambati, J., Ambati, B.K., Yoo, S.H., Ianchulev, S., and Adamis, A.P. (2003). Age-related macular degeneration: etiology, pathogenesis, and therapeutic strategies. Surv. Ophthalmol. 48, 257–293.

Andoniadou, C.L., Signore, M., Sajedi, E., Gaston-Massuet, C., Kelberman, D., Burns, A.J., Itasaki, N., Dattani, M., and Martinez-Barbera, J.P. (2007). Lack of the murine homeobox gene Hesx1 leads to a posterior transformation of the anterior forebrain. Development 134, 1499–1508.

Bartuma, H., Petrus-Reurer, S., Aronsson, M., Westman, S., André, H., and Kvanta, A. (2015). In Vivo Imaging of Subretinal Bleb-Induced Outer Retinal Degeneration in the Rabbit. Invest. Ophthalmol. Vis. Sci. 56, 2423–2430.

Begbie, J. (2013). Induction and patterning of neural crest and ectodermal placodes and their derivatives. Comprehensive Developmental Neuroscience: Patterning and Cell Type Specification in the Developing CNS and PNS 239–258.

Bergen, V., Lange, M., Peidli, S., Alexander Wolf, F., and Theis, FJ. (2019). Generalizing RNA velocity to transient cell states through dynamical modeling.

Bhatia, B., Singhal, S., Jayaram, H., Khaw, P.T., and Limb, G.A. (2010). Adult retinal stem cells revisited. Open Ophthalmol. J. 4, 30–38.

Blixt, Å., Mahlapuu, M., Aitola, M., Pelto-Huikko, M., Enerbäck, S., and Carlsson, P. (2000). A forkhead gene, FoxE3, is essential for lens epithelial proliferation and closure of the lens vesicle. Genes Dev. 14, 245–254.

Boles, N.C., Fernandes, M., Swigut, T., Srinivasan, R., Schiff, L., Rada-Iglesias, A., Wang, Q., Saini, J.S., Kiehl, T., Stern, J.H., et al. (2020). Epigenomic and Transcriptomic Changes During Human RPE EMT in a Stem Cell Model of Epiretinal Membrane Pathogenesis and Prevention by Nicotinamide. Stem Cell Reports 14, 631–647.

Bosze, B., Hufnagel, R.B., and Brown, N.L. (2020). Chapter 21 - Specification of retinal cell types. In Patterning and Cell Type Specification in the Developing CNS and PNS (Second Edition), J. Rubenstein, P. Rakic, B. Chen, and K.Y. Kwan, eds. (Academic Press), pp. 481–504.

Brandl, C., Zimmermann, S.J., Milenkovic, V.M., Rosendahl, S.M.G., Grassmann, F., Milenkovic, A., Hehr, U., Federlin, M., Wetzel, C.H., Helbig, H., et al. (2014). In-depth characterisation of Retinal Pigment Epithelium (RPE) cells derived from human induced pluripotent stem cells (hiPSC). Neuromolecular Med. 16, 551–564.

Butler, A., Hoffman, P., Smibert, P., Papalexi, E., and Satija, R. (2018). Integrating single-cell transcriptomic data across different conditions, technologies, and species. Nat. Biotechnol. 36, 411–420.

Cajal, M., Lawson, K.A., Hill, B., Moreau, A., Rao, J., Ross, A., Collignon, J., and Camus, A. (2012). Clonal and molecular analysis of the prospective anterior neural boundary in the mouse embryo. Development 139, 423–436.

Chen, J., Tambalo, M., Barembaum, M., Ranganathan, R., Simões-Costa, M., Bronner, M.E., and Streit, A. (2017). A systems-level approach reveals new gene regulatory modules in the developing ear. Development 144, 1531–1543.

Choudhary, P., and Whiting, P.J. (2016). A strategy to ensure safety of stem cell-derived retinal pigment epithelium cells. Stem Cell Res. Ther. 7, 127.

Cohen-Salmon, M., El-Amraoui, A., Leibovici, M., and Petit, C. (1997). Otogelin: a glycoprotein specific to the acellular membranes of the inner ear. Proc. Natl. Acad. Sci. U. S. A. 94, 14450–14455.

Collin, J., Queen, R., Zerti, D., Dorgau, B., Hussain, R., Coxhead, J., Cockell, S., and Lako, M. (2019). Deconstructing Retinal Organoids: Single Cell RNA-Seq Reveals the Cellular Components of Human Pluripotent Stem Cell-Derived Retina. Stem Cells 37, 593–598.

Cowan, C.S., Renner, M., De Gennaro, M., Gross-Scherf, B., Goldblum, D., Hou, Y., Munz, M., Rodrigues, T.M., Krol, J., Szikra, T., et al. (2020). Cell Types of the Human Retina and Its Organoids at Single-Cell Resolution. Cell 182, 1623–1640.e34.

Crespo-Enriquez, I., Partanen, J., Martinez, S., and Echevarria, D. (2012). Fgf8-Related Secondary Organizers Exert Different Polarizing Planar Instructions along the Mouse Anterior Neural Tube. PLoS One 7, e39977.

da Cruz, L., Fynes, K., Georgiadis, O., Kerby, J., Luo, Y.H., Ahmado, A., Vernon, A., Daniels, J.T., Nommiste, B., Hasan, S.M., et al. (2018). Phase 1 clinical study of an embryonic stem cell-derived retinal pigment epithelium patch in age-related macular degeneration. Nat. Biotechnol. 36, 328–337.

Cuomo, A.S.E., Seaton, D.D., McCarthy, D.J., Martinez, I., Bonder, M.J., Garcia-Bernardo, J., Amatya, S., Madrigal, P., Isaacson, A., Buettner, F., et al. (2020). Single-cell RNA-sequencing of differentiating iPS cells reveals dynamic genetic effects on gene expression. Nat. Commun. 11, 810.

Cvekl, A., and Wang, W.-L. (2009). Retinoic acid signaling in mammalian eye development. Exp. Eye Res. 89, 280–291.

Eagleson, G., Ferreiro, B., and Harris, W.A. (1995). Fate of the anterior neural ridge and the morphogenesis of the Xenopus forebrain. J. Neurobiol. 28, 146–158.

Frank, E., and Hall, M. (2001). A Simple Approach to Ordinal Classification. In Machine Learning: ECML 2001, (Springer Berlin Heidelberg), pp. 145–156.

Fuhrmann, S. (2010). Eye morphogenesis and patterning of the optic vesicle. Curr. Top. Dev. Biol. 93, 61–84.

Fuhrmann, S., Levine, E.M., and Reh, T.A. (2000). Extraocular mesenchyme patterns the optic vesicle during early eye development in the embryonic chick. Development 127, 4599–4609.

Fuhrmann, S., Zou, C., and Levine, E.M. (2014). Retinal pigment epithelium development, plasticity, and tissue homeostasis. Exp. Eye Res. 123, 141–150.

Fujimura, N. (2016). WNT/β-Catenin Signaling in Vertebrate Eye Development. Frontiers in Cell and Developmental Biology 4, 138.

Gao, Z., Mao, C.-A., Pan, P., Mu, X., and Klein, W.H. (2014). Transcriptome of Atoh7 retinal progenitor cells identifies new Atoh7-dependent regulatory genes for retinal ganglion cell formation. Dev. Neurobiol. 74, 1123–1140.

Gehrs, K.M., Anderson, D.H., Johnson, L.V., and Hageman, G.S. (2006). Age-related macular degeneration--emerging pathogenetic and therapeutic concepts. Ann. Med. 38, 450–471.

Gitton, Y., Benouaiche, L., Vincent, C., Heude, E., Soulika, M., Bouhali, K., Couly, G., and Levi, G. (2011). Dlx5 and Dlx6 expression in the anterior neural fold is essential for patterning the dorsal nasal capsule. Development 138, 897–903.

Grove, E.A., and Monuki, E.S. (2020). Chapter 1 - Morphogens, patterning centers, and their mechanisms of action. In Patterning and Cell Type Specification in the Developing CNS and PNS (Second Edition), J. Rubenstein, P. Rakic, B. Chen, and K.Y. Kwan, eds. (Academic Press), pp. 3–21.

Hastie, T., and Stuetzle, W. (1989). Principal Curves. J. Am. Stat. Assoc. 84, 502–516.

Hu, Y., Wang, X., Hu, B., Mao, Y., Chen, Y., Yan, L., Yong, J., Dong, J., Wei, Y., Wang, W., et al. (2019). Dissecting the transcriptome landscape of the human fetal neural retina and retinal pigment epithelium by single-cell RNA-seq analysis. PLoS Biol. 17, e3000365.

Imuta, Y., Nishioka, N., Kiyonari, H., and Sasaki, H. (2009). Short limbs, cleft palate, and delayed formation of flat proliferative chondrocytes in mice with targeted disruption of a putative protein kinase gene, Pkdcc (AW548124). Dev. Dyn. 238, 210–222.

Jung, A.R., Jung, C.-H., Noh, J.K., Lee, Y.C., and Eun, Y.-G. (2020). Epithelial-mesenchymal transition gene signature is associated with prognosis and tumor microenvironment in head and neck squamous cell carcinoma. Sci. Rep. 10, 3652.

Kagiyama, Y., Gotouda, N., Sakagami, K., Yasuda, K., Mochii, M., and Araki, M. (2005). Extraocular dorsal signal affects the developmental fate of the optic vesicle and patterns the optic neuroepithelium. Dev. Growth Differ. 47, 523–536.

Kasberg, A.D., Brunskill, E.W., and Steven Potter, S. (2013). SP8 regulates signaling centers during craniofacial development. Dev. Biol. 381, 312–323.

Kashani, A.H., Lebkowski, J.S., Rahhal, F.M., Avery, R.L., Salehi-Had, H., Dang, W., Lin, C.-M., Mitra, D., Zhu, D., Thomas, B.B., et al. (2018). A bioengineered retinal pigment epithelial monolayer for advanced, dry age-related macular degeneration. Sci. Transl. Med. 10.

Kulkarni, A., Anderson, A.G., Merullo, D.P., and Konopka, G. (2019). Beyond bulk: a review of single cell transcriptomics methodologies and applications. Curr. Opin. Biotechnol. 58, 129–136.

Kumamoto, T., and Hanashima, C. (2017). Evolutionary conservation and conversion of Foxg1 function in brain development. Dev. Growth Differ. 59, 258–269.

Kumar, P., Tan, Y., and Cahan, P. (2017). Understanding development and stem cells using single cell-based analyses of gene expression. Development 144, 17–32.

La Manno, G., Gyllborg, D., Codeluppi, S., Nishimura, K., Salto, C., Zeisel, A., Borm, L.E., Stott, S.R.W., Toledo, E.M., Villaescusa, J.C., et al. (2016). Molecular Diversity of Midbrain Development in Mouse, Human, and Stem Cells. Cell 167, 566–580.e19.

La Manno, G., Soldatov, R., Zeisel, A., Braun, E., Hochgerner, H., Petukhov, V., Lidschreiber, K., Kastriti, M.E., Lönnerberg, P., Furlan, A., et al. (2018). RNA velocity of single cells. Nature 560, 494–498.

La Manno, G., Siletti, K., Furlan, A., Gyllborg, D., and Vinsland, E. (2020). Molecular architecture of the developing mouse brain. BioRxiv.

Lederer, A.R., and La Manno, G. (2020). The emergence and promise of single-cell temporal-omics approaches. Curr. Opin. Biotechnol. 63, 70–78.

Lo Giudice, Q., Leleu, M., La Manno, G., and Fabre, P.J. (2019). Single-cell transcriptional logic of cell-fate specification and axon guidance in early-born retinal neurons. Development 146.

Lukowski, S.W., Lo, C.Y., Sharov, A.A., Nguyen, Q., Fang, L., Hung, S.S., Zhu, L., Zhang, T., Grunert, U., Nguyen, T., et al. (2019). A single-cell transcriptome atlas of the adult human retina. EMBO J. 38, e100811.

MacLean, A.L., Hong, T., and Nie, Q. (2018). Exploring intermediate cell states through the lens of single cells. Curr Opin Syst Biol 9, 32–41.

Mandai, M., Watanabe, A., Kurimoto, Y., Hirami, Y., Morinaga, C., Daimon, T., Fujihara, M., Akimaru, H., Sakai, N., Shibata, Y., et al. (2017). Autologous Induced Stem-Cell–Derived Retinal Cells for Macular Degeneration. N. Engl. J. Med. 376, 1038–1046.

Mao, X., An, Q., Xi, H., Yang, X.J., Zhang, X., Yuan, S., Wang, J., Hu, Y., Liu, Q., and Fan, G. (2019). Single-Cell RNA Sequencing of hESC-Derived 3D Retinal Organoids Reveals Novel Genes Regulating RPC Commitment in Early Human Retinogenesis. Stem Cell Reports 13, 747–760.

Marquardt, T., Ashery-Padan, R., Andrejewski, N., Scardigli, R., Guillemot, F., and Gruss, P. (2001). Pax6 is required for the multipotent state of retinal progenitor cells. Cell 105, 43–55.

Martínez-Morales, J.R., Dolez, V., Rodrigo, I., Zaccarini, R., Leconte, L., Bovolenta, P., and Saule, S. (2003). OTX2 activates the molecular network underlying retina pigment epithelium differentiation. J. Biol. Chem. 278, 21721–21731.

McCracken, I.R., Taylor, R.S., Kok, F.O., de la Cuesta, F., Dobie, R., Henderson, B.E.P., Mountford, J.C., Caudrillier, A., Henderson, N.C., Ponting, C.P., et al. (2020). Transcriptional dynamics of pluripotent stem cell-derived endothelial cell differentiation revealed by single-cell RNA sequencing. Eur. Heart J. 41, 1024–1036.

Menon, M., Mohammadi, S., Davila-Velderrain, J., Goods, B.A., Cadwell, T.D., Xing, Y., Stemmer-Rachamimov, A., Shalek, A.K., Love, J.C., Kellis, M., et al. (2019). Single-cell transcriptomic atlas of the human retina identifies cell types associated with age-related macular degeneration. Nat. Commun. 10, 4902.

Nishiguchi, S., Wood, H., and Kondoh, H. (1998). Sox1 directly regulates the γ-crystallin genes and is essential for lens development in mice. Genes.

Peng, G., and Westerfield, M. (2006). Lhx5 promotes forebrain development and activates transcription of secreted Wnt antagonists. Development 133, 3191–3200.

Petrus-Reurer, S., Bartuma, H., Aronsson, M., Westman, S., Lanner, F., André, H., and Kvanta, A. (2017). Integration of Subretinal Suspension Transplants of Human Embryonic Stem Cell-Derived Retinal Pigment Epithelial Cells in a Large-Eyed Model of Geographic Atrophy. Invest. Ophthalmol. Vis. Sci. 58, 1314–1322.

Petrus-Reurer, S., Bartuma, H., Aronsson, M., Westman, S., Lanner, F., and Kvanta, A. (2018). Subretinal Transplantation of Human Embryonic Stem Cell Derived-retinal Pigment Epithelial Cells into a Largeeyed Model of Geographic Atrophy. J. Vis. Exp.

Plaza Reyes, A., Petrus-Reurer, S., Antonsson, L., Stenfelt, S., Bartuma, H., Panula, S., Mader, T., Douagi, I., André, H., Hovatta, O., et al. (2016). Xeno-Free and Defined Human Embryonic Stem Cell-Derived Retinal Pigment Epithelial Cells Functionally Integrate in a Large-Eyed Preclinical Model. Stem Cell Reports 6, 9–17.

Plaza Reyes, A., Petrus-Reurer, S., Padrell Sanchez, S., Kumar, P., Douagi, I., Bartuma, H., Aronsson, M., Westman, S., Lardner, E., Andre, H., et al. (2020a). Identification of cell surface markers and establishment of monolayer differentiation to retinal pigment epithelial cells. Nat. Commun. 11, 1609.

Plaza Reyes, A., Petrus-Reurer, S., Sánchez, S.P., Kumar, P., Douagi, I., Bartuma, H., Aronsson, M., Westman, S., Lardner, E., Falk, A., et al. (2020b). Xeno-free, chemically defined and scalable monolayer differentiation protocol for retinal pigment epithelial cells.

Quadrato, G., Nguyen, T., Macosko, E.Z., Sherwood, J.L., Min Yang, S., Berger, D.R., Maria, N., Scholvin, J., Goldman, M., Kinney, J.P., et al. (2017). Cell diversity and network dynamics in photosensitive human brain organoids. Nature 545, 48–53.

Rheaume, B.A., Jereen, A., Bolisetty, M., Sajid, M.S., Yang, Y., Renna, K., Sun, L., Robson, P., and Trakhtenberg, E.F. (2018). Single cell transcriptome profiling of retinal ganglion cells identifies cellular subtypes. Nat. Commun. 9, 2759.

Rodin, S., Antonsson, L., Niaudet, C., Simonson, O.E., Salmela, E., Hansson, E.M., Domogatskaya, A., Xiao, Z., Damdimopoulou, P., Sheikhi, M., et al. (2014). Clonal culturing of human embryonic stem cells on laminin-521/E-cadherin matrix in defined and xeno-free environment. Nat. Commun. 5, 3195.

Saint-Jeannet, J.-P., and Moody, S.A. (2014). Establishing the pre-placodal region and breaking it into placodes with distinct identities. Dev. Biol. 389, 13–27.

Salero, E., Blenkinsop, T.A., Corneo, B., Harris, A., Rabin, D., Stern, J.H., and Temple, S. (2012). Adult human RPE can be activated into a multipotent stem cell that produces mesenchymal derivatives. Cell Stem Cell 10, 88–95.

Sarkar, A., Marchetto, M.C., and Gage, F.H. (2020). Chapter 9 - Neural induction of embryonic stem/induced pluripotent stem cells. In Patterning and Cell Type Specification in the Developing CNS and PNS (Second Edition), J. Rubenstein, P. Rakic, B. Chen, and K.Y. Kwan, eds. (Academic Press), pp. 185–203.

Satija, R., Farrell, J.A., Gennert, D., Schier, A.F., and Regev, A. (2015). Spatial reconstruction of single-cell gene expression data. Nat. Biotechnol. 33, 495–502.

Schmitt, S., Aftab, U., Jiang, C., Redenti, S., Klassen, H., Miljan, E., Sinden, J., and Young, M. (2009). Molecular characterization of human retinal progenitor cells. Invest. Ophthalmol. Vis. Sci. 50, 5901–5908.

Schwartz, S.D., Tan, G., Hosseini, H., and Nagiel, A. (2016). Subretinal Transplantation of Embryonic Stem Cell–Derived Retinal Pigment Epithelium for the Treatment of Macular Degeneration: An Assessment at 4 Years. Invest. Ophthalmol. Vis. Sci. 57, ORSFc1–ORSFc9.

Seo, S., Chen, L., Liu, W., Zhao, D., Schultz, K.M., Sasman, A., Liu, T., Zhang, H.F., Gage, P.J., and Kume, T. (2017). Foxc1 and Foxc2 in the Neural Crest Are Required for Ocular Anterior Segment Development. Invest. Ophthalmol. Vis. Sci. 58, 1368–1377.

Shao, J., Zhou, P.Y., and Peng, G.H. (2017). Experimental Study of the Biological Properties of Human Embryonic Stem Cell-Derived Retinal Progenitor Cells. Sci. Rep. 7, 42363.

Shekhar, K., Lapan, S.W., Whitney, I.E., Tran, N.M., Macosko, E.Z., Kowalczyk, M., Adiconis, X., Levin, J.Z., Nemesh, J., Goldman, M., et al. (2016). Comprehensive Classification of Retinal Bipolar Neurons by Single-Cell Transcriptomics. Cell 166, 1308–1323 e30.

Singh, S., and Groves, A.K. (2016). The molecular basis of craniofacial placode development. Wiley Interdiscip. Rev. Dev. Biol. 5, 363–376.

Soldatov, R., Kaucka, M., Kastriti, M.E., and Petersen, J. (2019). Spatiotemporal structure of cell fate decisions in murine neural crest.

Song, W.K., Park, K.-M., Kim, H.-J., Lee, J.H., Choi, J., Chong, S.Y., Shim, S.H., Del Priore, L.V., and Lanza, R. (2015). Treatment of macular degeneration using embryonic stem cell-derived retinal pigment epithelium: preliminary results in Asian patients. Stem Cell Reports 4, 860–872.

Sparrow, J.R., Hicks, D., and Hamel, C.P. (2010). The retinal pigment epithelium in health and disease. Curr. Mol. Med. 10, 802–823.

Sridhar, A., Hoshino, A., Finkbeiner, C.R., Chitsazan, A., Dai, L., Haugan, A.K., Eschenbacher, K.M., Jackson, D.L., Trapnell, C., Bermingham-McDonogh, O., et al. (2020). Single-Cell Transcriptomic Comparison of Human Fetal Retina, hPSC-Derived Retinal Organoids, and Long-Term Retinal Cultures. Cell Rep. 30, 1644–1659 e4.

Streit, A. (2007). The preplacodal region: an ectodermal domain with multipotential progenitors that contribute to sense organs and cranial sensory ganglia. Int. J. Dev. Biol. 51, 447–461.

Sunness, J.S. (1999). The natural history of geographic atrophy, the advanced atrophic form of age-related macular degeneration. Mol. Vis. 5, 25.

Tahayato, A., Dollé, P., and Petkovich, M. (2003). Cyp26C1 encodes a novel retinoic acid-metabolizing enzyme expressed in the hindbrain, inner ear, first branchial arch and tooth buds during murine development. Gene Expr. Patterns 3, 449–454.

Velasco, S., Kedaigle, A.J., Simmons, S.K., Nash, A., Rocha, M., Quadrato, G., Paulsen, B., Nguyen, L., Adiconis, X., Regev, A., et al. (2019). Individual brain organoids reproducibly form cell diversity of the human cerebral cortex. Nature 570, 523–527.

Veres, A., Faust, A.L., Bushnell, H.L., Engquist, E.N., Kenty, J.H.-R., Harb, G., Poh, Y.-C., Sintov, E., Gürtler, M., Pagliuca, F.W., et al. (2019). Charting cellular identity during human in vitro β-cell differentiation. Nature 569, 368–373.

Voigt, A.P., Mulfaul, K., Mullin, N.K., Flamme-Wiese, M.J., Giacalone, J.C., Stone, E.M., Tucker, B.A., Scheetz, T.E., and Mullins, R.F. (2019). Single-cell transcriptomics of the human retinal pigment epithelium and choroid in health and macular degeneration. Proc. Natl. Acad. Sci. U. S. A. 116, 24100–24107.

Wang, Y., Tang, Z., and Gu, P. (2020). Stem/progenitor cell-based transplantation for retinal degeneration: a review of clinical trials. Cell Death Dis. 11, 793.

Wolf, F.A., Angerer, P., and Theis, F.J. (2018). SCANPY: large-scale single-cell gene expression data analysis. Genome Biol. 19, 15.

Yamada, R., Mizutani-Koseki, Y., Hasegawa, T., Osumi, N., Koseki, H., and Takahashi, N. (2003). Cell-autonomous involvement of Mab21l1 is essential for lens placode development. Development 130, 1759–1770.

Yun, S., Saijoh, Y., Hirokawa, K.E., Kopinke, D., Murtaugh, L.C., Monuki, E.S., and Levine, E.M. (2009). Lhx2 links the intrinsic and extrinsic factors that control optic cup formation. Development 136, 3895–3906.

Zarbin, M., Sugino, I., and Townes-Anderson, E. (2019). Concise review: update on retinal pigment epithelium transplantation for age-related macular degeneration. Stem Cells Transl. Med. 8, 466–477.

Zeisel, A., Hochgerner, H., Lönnerberg, P., Johnsson, A., Memic, F., van der Zwan, J., Häring, M., Braun, E., Borm, L.E., La Manno, G., et al. (2018). Molecular Architecture of the Mouse Nervous System. Cell 174, 999–1014.e22.

Zhou, J., Wang, C., Wang, Z., Dampier, W., Wu, K., Casimiro, M.C., Chepelev, I., Popov, V.M., Quong, A., Tozeren, A., et al. (2010). Attenuation of Forkhead signaling by the retinal determination factor DACH1. Proc. Natl. Acad. Sci. U. S. A. 107, 6864–6869.

