## Supplementary material for "Molecular profiling of retinal pigment epithelial cell differentiation for therapeutic use": Table S1

**Table S1. Summary of the single-cell RNA-sequencing datasets obtained for the characterization of hESC-RPE differentiation protocol and the transcriptomic landscape of the retina.** Additional datasets are obtained from the literature and are used and cited appropriately.

| **Dataset Name** | **Dataset Description** | **Approximate Number of Cells (after QC filtering)** | **Source** |
| --- | --- | --- | --- |
| ***In vitro***  **hESC-RPE differentiation**  **60-day time course** | Seven individual time points: days 7, 14, 30, 38, 45, 60, and undifferentiated hESC control | 11,791 cells | Lanner Lab  (KI) |
| **Embryonic optic vesicle** | Carnegie Stage 13 (about week 5) | 2,637 cells | Kriegstein Lab (UCSF) |
| **Human fetal eyes** | Two fetal eyes (week 7.5) | 2,742 cells | Lanner Lab  (KI) |
| ***In vitro* NCAM1-High sorted hESC-RPE D30 + altered differentiation** | Using altered protocol for 40 days  (Shao et al, 2017) | 980 cells | Lanner Lab  (KI) |
| ***In vivo* transplanted hESC-RPE** | D60 hESC-RPE subretinally injected into albino rabbit eye for 30 days | 65 cells | Lanner Lab  (KI) |
| **Albino rabbit eyes** | Sequenced jointly with *in vivo* transplanted hESC-RPEs | 1,965 cells | Lanner Lab  (KI) |
| **Human adult eyes** | Two adult eyes  (45 years old) | 5,538 cells | Lanner Lab  (KI) |
